# Integrated Respirometry and Metabolomics Unveil Circadian Metabolic Dynamics in Drosophila

**DOI:** 10.1101/2025.07.22.665580

**Authors:** Farheen Akhtar, Dania Malik, Arjun Sengupta, Paula Haynes, Andrew D. Nguyen, Jaco Klok, Amita Sehgal, Aalim M. Weljie

## Abstract

Sleep and circadian rhythms shape organismal energy patterns, but how this timing connects to oxygen use and carbon dioxide production remains incompletely understood. We combined high-resolution respirometry with LC-MS-based metabolomics to characterize respiratory dynamics and metabolic states in *Drosophila melanogaster*, resolving genotype-specific impacts of sleep disruption and circadian regulation. Wild-type flies under light-dark cycles (WT-LD) showed rhythmic respiratory patterns reflective of anticipatory coordination of mitochondrial energy metabolism, amino acid turnover, and redox cycling. Short-sleep mutants (*fmn*, *sss*) exhibited elevated metabolic rates with reactive shifts of fuel preferences toward lipid and amino acid catabolism and altered mitochondrial respiration. The clock-mutant (*per^01^*) and flies under constant darkness (WT-DD) showed reactive and widespread metabolic dysregulation and impaired redox homeostasis. These findings demonstrate that both sleep and circadian systems contribute to aligning metabolic substrate selection with energy demands, offering mechanistic insights into how disruptions in behavioral states compromise metabolic health.

## INTRODUCTION

Metabolism is a fundamental physiological process responsible for coordinating energy production and substrate utilization in response to changing physiological demands. This process is dynamically regulated by behavioral states, such as sleep and wakefulness, as well as directly by internal circadian clocks that align physiology with the external environment [1-3]. Disruptions in these temporal processes are strongly associated with metabolic disorders, including obesity and type 2 diabetes [4-6] as well as other metabolic diseases such as cardiovascular dysfunction and cancer [7].

With conserved and well-characterized sleep and circadian mechanisms [8, 9], as well as metabolic pathways also found in mammals, *Drosophila* offers a genetically tractable model for dissecting interactions between these different systems. In particular, sleep mutants with robust phenotypes can be employed to determine how loss of sleep impacts metabolic parameters, including rhythms of metabolic activity. At the same time, additional tools are now available for metabolic measurements previously not possible. Multiple approaches can be combined for a comprehensive understanding of metabolism under precisely controlled environmental conditions and at high temporal resolution [10].

Indirect calorimetry is a non-invasive form of respirometry that enables real-time quantification of whole-organism metabolic rate by measuring oxygen consumption (VO₂) and carbon dioxide production (VCO₂)[11, 12]. These measurements allow for the estimation of energy expenditure and respiratory quotient (RQ), which can provide an index of relative substrate utilization, with appropriate caveats [13, 14]. Although widely applied in mammalian systems [15], indirect calorimetry in small model organisms such as *Drosophila melanogaster* has only recently become feasible with advances in high-resolution, small-volume flow-through systems [16-18]. This is a critical advancement, as dissecting time-of-day-dependent metabolic regulation in mammals is often confounded by overlapping effects of sleep, activity, and feeding rhythms [19, 20]. While this can also be true in flies, the genetic tools available allow distinction between these processes to a greater extent.

This study focuses on wild-type isogenic control flies alongside three genetically characterized mutants: two short-sleep mutants: *fumin* (*fmn*), which displays impaired dopamine transporter function and defective dopamine reuptake [21] and *sleepless* (*sss*), which lacks the Sleepless protein, resulting in altered membrane excitability and reduced GABA signaling [22-24]; and the circadian mutant *period^01^* (*per^01^*), which lacks a functional molecular clock [25]. To investigate how sleep and circadian disruption independently and interactively influence metabolic regulation, we utilized a small-animal respirometry platform built on the MAVEn-FT system, coupled with a Licor 7000 CO₂ analyzer and an Oxzilla differential O₂ analyzer. This system builds upon the Sleep and Activity Metabolic Monitor (SAMM), which first demonstrated the feasibility of simultaneous CO₂ and O₂ measurements in *Drosophila* [26]. This MAVEn-based platform enhances temporal resolution and throughput, enabling continuous measurement of respiratory parameters across multiple groups of flies over daily cycles, and allowing fine-scale profiling of metabolic rhythms and their disruption across sleep-wake cycles. In this study, we interpret RQ conservatively as reflecting relative shifts in substrate utilization over time and between genotypes, rather than as a precise quantitative measure of absolute carbohydrate versus lipid oxidation.

To provide a more comprehensive perspective on metabolic regulation, we complemented newly generated respiratory measurements with steady-state metabolomic profiling using liquid chromatography-mass spectrometry (LC-MS) data previously published from our group [27]. This integrative framework enabled time-matched and lag-aware analysis of respiratory output and metabolite profiles across Zeitgeber time in the LD cycle, providing detailed insights into how sleep and circadian mutations influence metabolic homeostasis [27-29]. By employing this framework, we sought to uncover conserved mechanisms linking temporal biological processes, such as sleep and circadian rhythms, to metabolic homeostasis and to elucidate how genetic and environmental perturbations in these systems influence energy balance and metabolic regulation.

## METHODS

### Drosophila Strains, Entrainment and Collection

*Drosophila melanogaster* isogenic control (*iso*), two short-sleep mutants, fumin (*fmn*) [21] and sleepless (*sss*) [22-24], as well as a circadian clock mutant (*per^01^*) [25]. were used for this study. Male flies were collected shortly after eclosion and entrained in light-dark (LD) incubators for a minimum of three days before diurnal (LD) time-course collection across Zeitgeber time (ZT). At the time of collection, flies were between five and seven days old. Wild-type (WT) isogenic control flies were either maintained under 12 hrs:12 hrs light versus dark conditions (WT-LD) or placed in constant darkness (WT-DD) for at least 24 hours prior to collection to examine light-independent rhythms. Mutant strains (*fmn, sss, and per^01^*) were maintained under LD cycles. All flies were reared on a standard cornmeal/molasses medium at 25°C. Zeitgeber time (ZT) 0 was designated as lights-on, with lights-off occurring at ZT12 (12/12 LD).

Only male flies were used for all respirometry experiments and for integration with the metabolomics dataset. This was done to reduce variability introduced by female reproductive status (e.g., mating/egg production) and associated metabolic differences, enabling a clearer comparison across genotypes and lighting conditions.

### Respirometry Setup

All experiments were performed using groups of 25 flies per chamber. To account for genotype-specific differences in body size, each group of 25 flies (*WT, fmn, sss, and per^01^*) was weighed prior to respirometry, and these weights were used to normalize VCO₂ and VO₂ values for accurate comparison. Each chamber was provisioned with 2 mL of fly food (2% agar, 5% sucrose) to sustain the flies during the 24-hour continuous recording period. Control chambers containing only food, without flies, were included to account for background microbial metabolism. To simulate natural microbial inoculation, flies were briefly introduced into these control chambers and then removed before data acquisition began. Each genotype measurement represents an average of ∼300 flies (25 flies/chamber × 4 chambers/experiment × 3 experimental days), with the chamber as the experimental unit.

The respirometry system continuously measured CO₂ production and O₂ consumption using a flow-through MAVEn system. It consisted of five main components: (1) a zero-grade air source and scrubbing column to remove residual CO₂, (2) mass-flow controllers to regulate airflow, (3) Sable Systems RC chambers to house the flies, (4) the MAVEn system to direct air sequentially from each chamber, and (5) gas analyzers for measuring CO₂ and O₂ concentrations. The system was calibrated every 3-5 trials using 100% N₂ (CO₂ = 0 ppm) and a known CO₂ standard to ensure measurement accuracy.

#### Zero-Grade Air Supply and Scrubbing System

Zero-grade compressed air (<0.1 ppm hydrocarbons) was first passed through a custom-built scrubbing column (#26800, Drierite) to remove residual CO₂. The column was packed with an inner layer of Ascarite (#81133–20–2, Acros Organics) flanked by two outer layers of Drierite (#7779–18–9, Drierite), separated by glass wool (#11–388, Fisher Scientific).

To re-humidify the air to ∼9 parts per thousand (ppt) water vapor content, it was then passed through Nafion tubing (#TT-070, Perma Pure, LLC) submerged in deionized water. Bev-A-line nonpermeable tubing (#56280, United States Plastic Corp.) was used to connect all system components.

#### Airflow Regulation

Re-humidified, CO₂-free air was split into two streams, each regulated by mass flow controllers to maintain a constant flow rate of 25 mL/min. One stream provided reference air to the CO₂ and O_2_ analyzers (controlled by a Side-Trak 840 Series, Sierra Instruments, Inc. MFCV), while the second stream provided flow to the fly chambers (controlled by the MAVEn’s internal mass flow-controlled system).

#### Respirometry Chambers and MAVEn System’s Airflow Management

Flies were housed in Sable Systems RC chambers (70 mm × 20 mm borosilicate glass tubes), integrated into a continuous flow through MAVEn system. The MAVEn splits airflow into 16 channels for fly chambers and one dedicated baseline channel. Airflow is constantly and continuously maintained through all fly chambers while the MAVEn’s multiplexing system will switch the ‘active’ chamber’s airflow toward the analyzer chain for a preset dwell time of 120 seconds per chamber, and 60 seconds for baseline, with a baseline interleave ratio of 4 to ensure accuracy in O₂ measurement (i.e a baseline measurement after every four chambers). In this system the MAVEn manages air flow to allow for the sequential measurements of individual chamber gas parameters and air flow rates. Experimental respirometry recording were conducted for 24-hour periods. System control and data acquisition were handled via Sable Systems MAVEn Controller software.

#### CO₂ Production Measurement

Baseline CO₂ levels were recorded from the reference air prior to entering the chambers. As flies respired, the CO₂ released in each chamber was subsequently flushed to the CO₂ analyzer (Li-7000, LI-COR Biosciences) via the MAVEn system.

#### O₂ Consumption Measurement

Similarly, O_2_ consumed by flies was measured immediately using the Oxzilla differential O_2_ analyzer (Oxzilla II Oxygen Analyzer, Sable Systems International) placed downstream from the CO_2_ analyzer in the analyzer chain. In this setup-controlled flow reference air exiting the CO_2_ analyzer’s reference cell was routed into the Oxzilla’s reference channel, while animal air exiting the CO_2_ analyzer’s sample cell was routed to the Oxzilla’s sample channel. To prevent dilution effects, water vapor was removed prior to O₂ analysis using small scrubbing columns.

#### Water Vapor Scrubbing Columns

Separate scrubbing columns were used for reference and animal airflows and placed in-line directly prior to the Oxzilla differential O_2_ analyzer’s reference and sample channel inlet ports. Each column was made from a 10 mL syringe body (#14955459, Fisher Scientific) filled with an inner layer of Ascarite and two outer layers of magnesium perchlorate (M54, Fisher Scientific), separated by glass wool. Bev-A-line tubing was secured with rubber stoppers (#14–135E, Fisher Scientific). Columns were replaced twice during each 24-hour experiment to maintain performance.

### Carbon Dioxide and Oxygen Analysis and Calculations

O₂ consumption was quantified by measuring the decrease in O₂ concentration as air passed through chambers containing flies. To ensure accuracy, a Savitzky-Golay filter (25-second window) was applied to smooth the raw signal, followed by a lag correction of 127 seconds accounted for delay between chamber exit and O_2_ analyzer input. To match dry atmospheric air, the reference air was baseline corrected to 20.95% O₂. For both reference and animal air the %O_2_ values were converted to O_2_ fractional contents by dividing percentage values by 100 (thus baseline air has a fractional content of 0.2095). The VO₂ value from an empty chamber was subtracted from experimental values [30]:

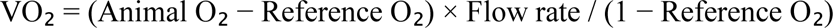

VO₂ values were expressed in μL/hr by summing all 5-minute bins into hourly averages.

CO_2_ production was quantified by measuring the increase in CO₂ concentration as air passed through chambers containing flies. To ensure accuracy, a Savitzky-Golay filter (25-second window) was applied to smooth the raw signal, followed by a lag correction of 22 seconds with Z correction and 28 seconds without Z correction accounted for delay between chamber exit and O_2_ analyzer input. To match dry atmospheric air, the reference air was baseline corrected to 0 parts per million (ppm) CO₂. For both reference and animal air the ppm CO_2_ values were converted to CO_2_ fractional contents by dividing percentage values by 1000000. The VCO₂ value from an empty chamber was subtracted from experimental values [30]:

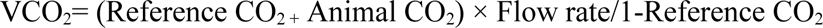

VCO₂ values were expressed in μL/hr by summing all 5-minute bins into hourly averages.

Respiratory quotient (RQ) was calculated as:

RQ= VCO_2_ / VO_2_. RQ was used as an index of relative shifts in substrate utilization over time and between genotypes, interpreted conservatively rather than as a precise quantitative measure of absolute carbohydrate versus lipid oxidation as this has not been directly characterized in flies.

### Gut Tissue Respirometry

Oxygen consumption rate (OCR) was measured in dissected gut tissue using the Resipher System (Lucid Scientific, GA, USA). Male flies were briefly anesthetized on ice and whole guts were dissected in Schneider’s Media. 96-well flat-bottom plates were coated with 50 µL of Poly-D-lysine for 30 min to promote tissue adhesion; each well was then filled with 300 µL of Schneider’s Media, a single gut was placed in a fixed position within each well, and the plate was fitted with the Resipher sensor lid. The Resipher hub recorded OCR (fmol/mm²/s) continuously for 72 h in a humidified incubator at 27 °C. OCR was extracted in 4-h bins and averaged over the 24-h window beginning 12 h after loading (the 12-36 h interval, once tissue respiration had stabilized). Experiments were performed in three independent runs; values were pooled, median-normalized to the *iso^31^* controls within each run, and log2(x + 10)-transformed. A single *iso^31^* gut in the third run of the *per^01^* experiment was identified as an outlier (Prism “Identify Outliers”) and excluded; no other values were removed. Normalized OCR was compared between each mutant and *iso^31^* using the Mann-Whitney test in GraphPad Prism 10, with significance at p < 0.05 (*fmn* vs *iso^31^*, n = 12 vs 12 guts; *per^01^* vs *iso^31^*, n = 12 vs 11 guts after the single *iso^31^* exclusion).

### Metabolite Extraction and LCMS Measurements

The metabolomics dataset analyzed in this study was previously published and is publicly available, as described in detail in [27, 31]. In the present study, these data were integrated with respirometry measurements to assess temporal relationships between metabolite abundance and respiratory output. Briefly, polar metabolites were extracted from fly bodies sampled every 2 hours from ZT0 to ZT22 using a modified Bligh-Dyer extraction method, as previously reported [31, 32]. The polar fraction of each extract was dried under vacuum and reconstituted in 100 μL of acetonitrile:Milli-Q water, followed by vortexing for 20 seconds. Samples were analyzed using an ion-switching LC-MS method, and peak integration and data processing were performed according to previously published procedures [27].

### Statistical Analysis

All statistical analyses were conducted using GraphPad Prism. Non-parametric Spearman correlation was used to evaluate relationships among metabolic parameters. Metabolites with *p*-values below 0.05 were considered statistically significant and selected for further investigation. To assess genotype-specific differences, one-way ANOVA was performed followed by a pairwise unpaired two-tailed Student’s *t*-tests comparing the control group (WT-LD) with each genotype (*fmn*, *sss*, *per^01^*, and WT-DD).

### Metabolomics-respirometry integration and lag analysis

To integrate respirometry with metabolomics, RQ was recorded continuously at 1-second resolution and averaged into 5-minute bins. Because steady-state metabolite measurements were acquired at 2-hour intervals, we extracted the RQ values corresponding to each 2-hour Zeitgeber Time (ZT) sampling point from the 5-minute-binned dataset to generate time-matched RQ-metabolite pairs. We additionally evaluated temporal relationships using a lag analysis by systematically shifting the RQ time series relative to the metabolite timepoints (−120, −60, −30, −15, −5, +5, +15, +30, +60, and +120 minutes). Metabolite abundances were normalized as described above, and associations between RQ and individual metabolites were quantified using Spearman rank correlations (ρ) at each lag. Metabolites showing strong associations (e.g., |ρ| > 0.7 with nominal p < 0.05) were carried forward for visualization and summary, and lag direction was interpreted as metabolites preceding (negative lag) or following (positive lag) changes in RQ.

### Rhythmicity analysis

Rhythmicity analysis was performed using Nitecap, a tool for circadian and rhythmic analyses [33]. Time-series metabolic data were analyzed to compute rhythmic parameters using the RAIN algorithm [34]. Rhythms were assessed for statistical significance using false discovery rate (FDR)-adjusted p-values, ensuring robust detection while minimizing false positives. The lag parameters of each rhythm were computed using JTK cycle algorithm [35].

### Pathway Analysis

Significant metabolites identified from univariate analyses were used for metabolic pathway analysis via MetaboAnalyst 5.0. Metabolites were uploaded using HMDB identifiers and analyzed using the hypergeometric enrichment method, with relative-betweenness centrality applied for topology analysis based on the *Drosophila melanogaster* (KEGG) pathway library. Pathways with a p-value less than 0.05 were considered statistically significant.

## RESULTS

### Metabolic Fuel Utilization Across Genotypes Based on Respiratory Quotient (RQ)

To assess whole-body metabolic activity, we performed respirometry on groups of 25 flies per genotype, measuring oxygen consumption (VO₂) and carbon dioxide production (VCO₂) across the 24-hour cycle. The respiratory quotient (RQ), calculated as the ratio of VCO₂ to VO₂, provides insight into the predominant metabolic substrate being utilized: an RQ of 1.0 reflects pure carbohydrate metabolism, values below 1.0 indicate lipid and protein oxidation, and values above 1.0 suggest lipogenesis (the conversion of carbohydrates into fats) [14]. RQ, VCO₂, and VO₂ were continuously recorded from Zeitgeber Time (ZT) 0 to 24 at one-second intervals and subsequently averaged into 5-minute bins for analysis. **Figures 1a-c** shows the 5-minute binned VCO₂, VO₂ and RQ profiles across the 24-hour cycle for all genotypes (separate individual plots for VCO₂, VO₂, and RQ for each genotype are provided in **Figure S1. a-e**, **S2. a-e and S3. a-e** respectively. Replicate information for respirometry provided in **Supplementary Table S3**). The average values of VCO₂, VO₂ and RQ across the full 24-hour cycle are summarized in **Figure 1d-f**.

**Figure 1.**
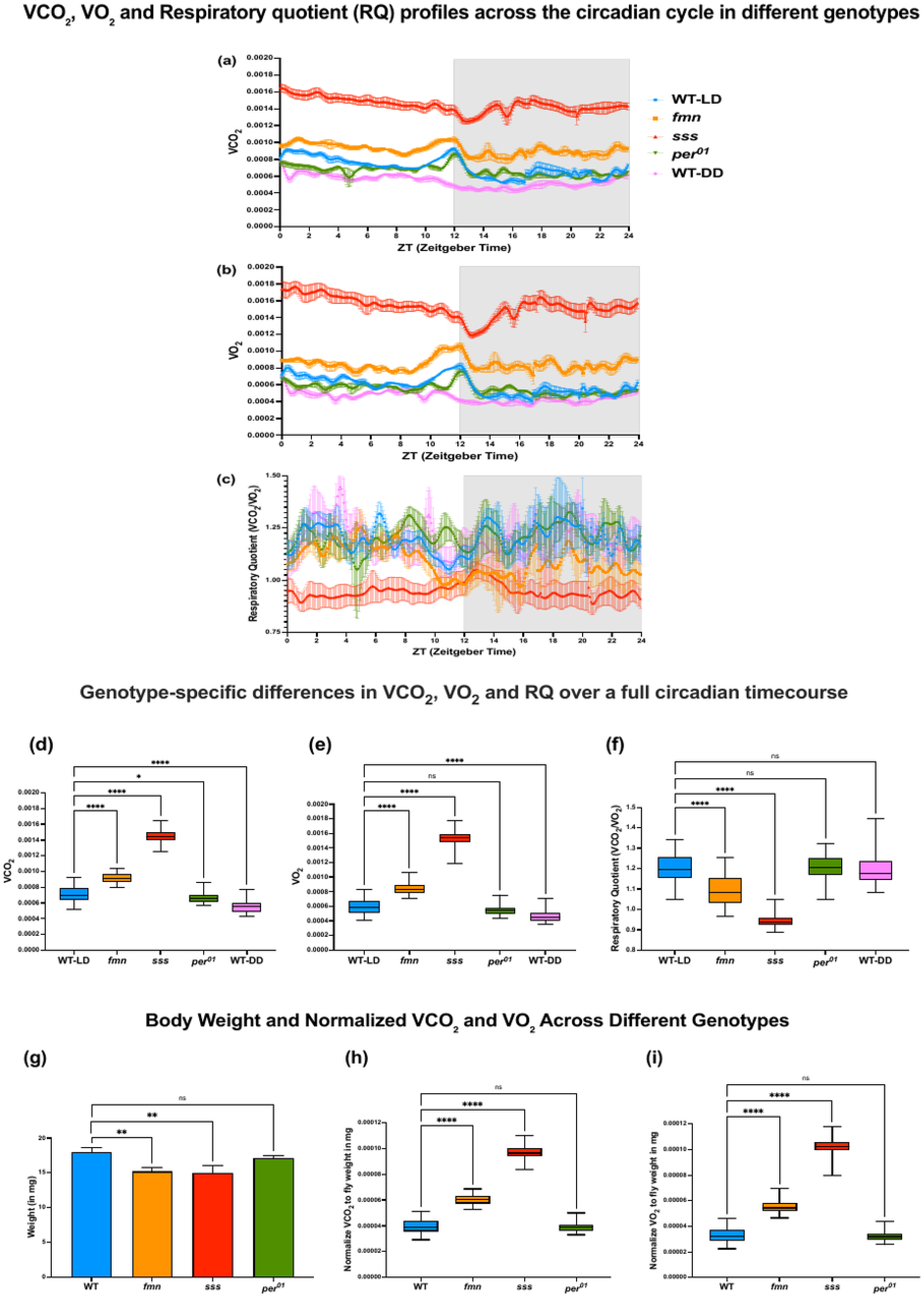
**a-c. *VCO₂, VO₂, and respiratory quotient (RQ) profiles across the 24-hour cycle in different genotypes*.** VCO₂, VO₂ and RQ (VCO₂/VO₂) was continuously recorded at one-second intervals from Zeitgeber Time (ZT) 0 to 24 and subsequently averaged into 5-minute bins for analysis. Traces represent mean VCO₂, VO₂, and RQ values across the 24-hour recording period for wild-type flies under light-dark conditions (WT-LD), short-sleep mutants *fumin* (*fmn*) and *sleepless* (*sss*), the circadian clock mutant *period*^01^ (*per*^01^), and wild-type flies maintained in constant darkness (WT-DD). Source data are the continuous respirometry recordings. **d-f. *Genotype-specific differences in VCO₂, VO₂, and RQ measured over a full 24-hour recording period*.** Boxplots show average values of (left to right) respiratory quotient (RQ), carbon dioxide production (VCO₂), and oxygen consumption (VO₂) across genotypes and lighting conditions. Measurements were taken continuously over a 24-hour period using a flow-through MAVEn system. Genotypes include wild-type under light-dark (WT-LD) and constant darkness (WT-DD), short-sleep mutants *fumin* (*fmn*) and *sleepless* (*sss*), and circadian mutant *per^01^*. Group differences were assessed using the **Kruskal-Wallis test**, followed by **Dunn’s multiple comparisons post hoc test**. Significance is denoted as: *p* < 0.05 (\**), p < 0.01 (*\*\**), p < 0.001 (*\*\*\*); ns = not significant. **g-i. *Body Weight and Normalized VCO₂ and VO₂ Across Different Genotypes*.** Body weight (mg) and respiratory parameters (VCO₂ and VO₂) normalized to body weight (mg) were measured in wild type (WT), *fmn*, *sss*, and *per^01^* mutant flies. Statistical significance was assessed using one-way ANOVA followed by Dunnett’s multiple comparisons test against WT. *p* < 0.05 (\**), p < 0.01 (*\*\**), p < 0.001 (*\*\*\*); ns = not significant. For all panels with error bars or shaded error bands, values are presented as mean ± SEM. Data represent approximately 300 flies per genotype across three experimental days (25 flies/chamber, four chambers/experiment). The chamber was used as the experimental unit; *n* denotes the number of chambers.

Among the genotypes tested, WT-LD, WT-DD, and *per^01^*flies exhibited similarly elevated RQ values (1.20, 1.19, and 1.20, respectively), consistent with active lipogenesis. Group differences were assessed using the Kruskal-Wallis test, followed by Dunn’s multiple comparisons post hoc test, which revealed no statistically significant differences in RQ among these three genotypes (**Figure 1f, ns**), suggesting that neither free-running conditions in constant darkness (WT-DD) nor loss of clock function in the *per^01^*mutant alters the dominant fuel utilization strategy under basal conditions.

In contrast, short-sleeping mutants displayed lower RQ values. The *fmn* mutant exhibited a moderate reduction (RQ = 1.09), reflecting increased reliance on carbohydrate metabolism. The *sss* mutant showed the lowest RQ (0.94), indicating a shift toward lipid and protein catabolism. Both mutants were significantly different from WT-LD (*p* < 0.001), highlighting genotype-specific alterations in fuel utilization associated with sleep loss (**Figure 1f**).

These results demonstrate that WT-LD, WT-DD, and *per^01^* flies maintain a lipogenic profile, whereas sleep-deficient mutants exhibit altered substrate utilization, favoring catabolism of carbohydrates or lipids depending on the severity of sleep disruption.

Furthermore, to account for potential differences in body size that could confound VCO₂ and VO₂ measurements, fly weights were recorded for each genotype before respirometry. Notably, *fmn* and *sss* mutants weighed significantly less than WT controls, whereas *per^01^* flies had comparable weights (**Figure 1g**). Accordingly, VCO₂ and VO₂ values were normalized to fly weight, and the genotype-specific differences remained robust after normalization (**Figure 1h-i**). These data confirm that the observed metabolic differences are not attributable to body size but reflect inherent alterations in metabolic fuel utilization. Figures 1, 2, 3, and Figure S4 are derived from the same underlying continuous respirometry recordings but present different analytical summaries: Figure 1 shows 5-minute-binned time-course traces and 24-hour averages, Figures 2-3 present rhythmicity and phase analyses, and Figure S4 presents day (ZT0-12) versus night (ZT12-24) averages.

**Figure 2.**
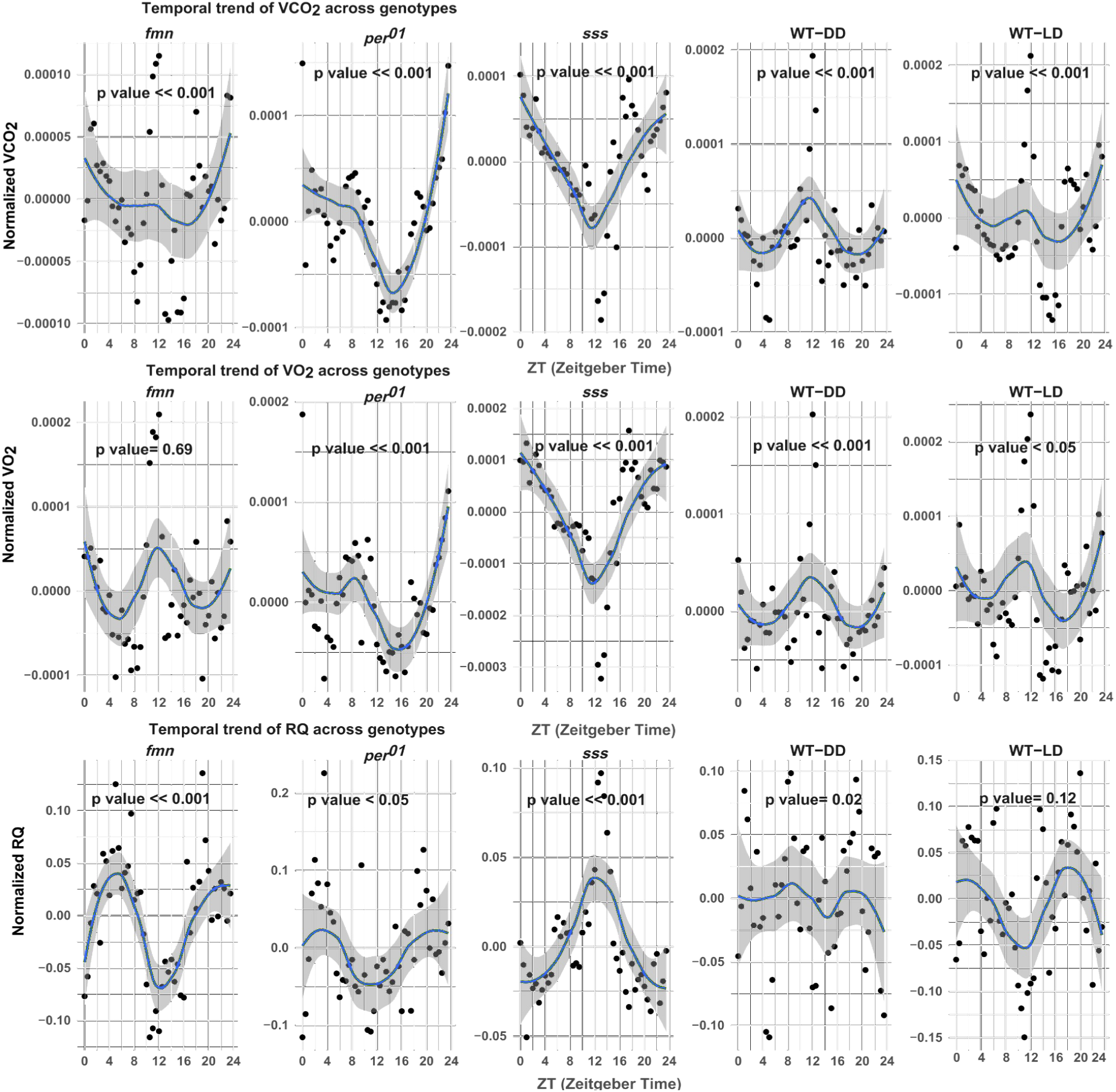
Twenty-four-hour time course of respiratory parameters across genotypes. Normalized temporal profiles of carbon dioxide production (VCO₂) (top), oxygen consumption (VO₂) (middle), and respiratory quotient (RQ) (bottom) plotted over a 24-hour cycle (ZT0–24) for genotypes: wild-type in light-dark (WT-LD) and constant darkness (WT-DD), short-sleep mutants *fumin* (*fmn*) and *sleepless* (*sss*), and circadian mutant *per^01^*. Curves reveal genotype-specific rhythmicity and metabolic dynamics across the 24-hour recording period. Significant genotype × time interactions were observed for all variables (VCO₂, VO₂, RQ), indicating genotype-dependent alterations in respiratory rhythmicity. *p* < 0.05 considered significant. For all panels with error bars or shaded error bands, values are presented as mean ± SEM. Source data are the same continuous respirometry recordings shown in Figure 1; Figure 2 presents rhythmicity analysis of VCO₂, VO₂, and RQ across the 24-hour cycle. Data represent approximately 300 flies per genotype across three experimental days (25 flies/chamber, four chambers/experiment). The chamber was used as the experimental unit; *n* denotes the number of chambers.

**Figure 3.**
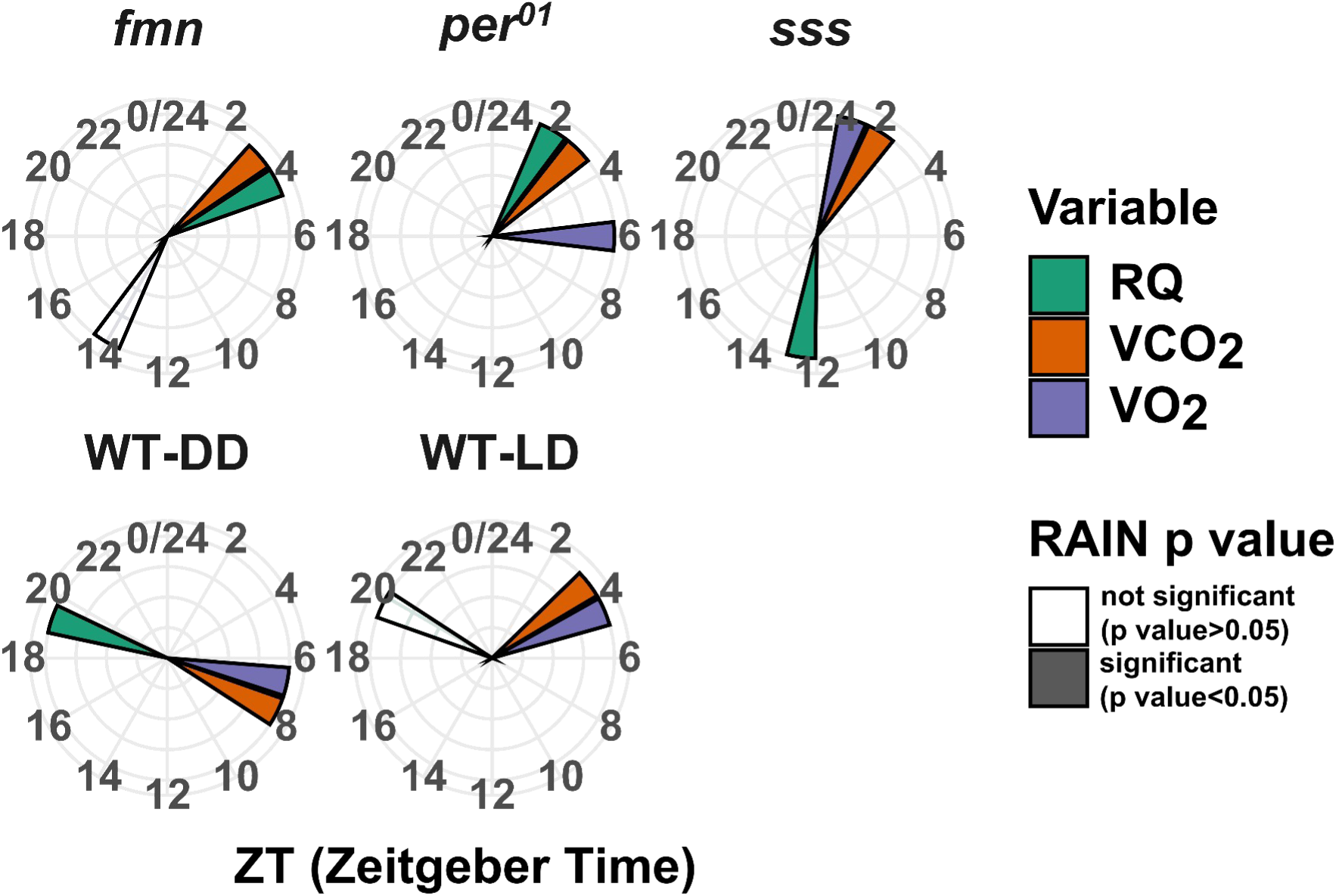
Phase distribution of respiratory rhythms across genotypes. Polar plots depicting the phase (peak timing) of rhythmic expression for carbon dioxide production (VCO₂), oxygen consumption (VO₂), and respiratory quotient (RQ) over a 24-hour cycle (ZT0-24) for each genotype. Genotypes include wild-type in light-dark (WT-LD) and constant darkness (WT-DD), short-sleep mutants *fumin* (*fmn*) and *sleepless* (*sss*), and circadian mutant *per^01^.* Statistical significance of rhythmicity was determined using the RAIN algorithm: darker-colored bars indicate significant rhythms (p < 0.05), and lighter-colored bars indicate non-significant rhythms (p > 0.05). Where applicable, plotted values are presented as mean ± SEM. Source data are the same continuous respirometry recordings shown in Figure 1; Figure 3 presents phase/peak-timing analysis derived from the rhythmicity analysis

**Table 1.**
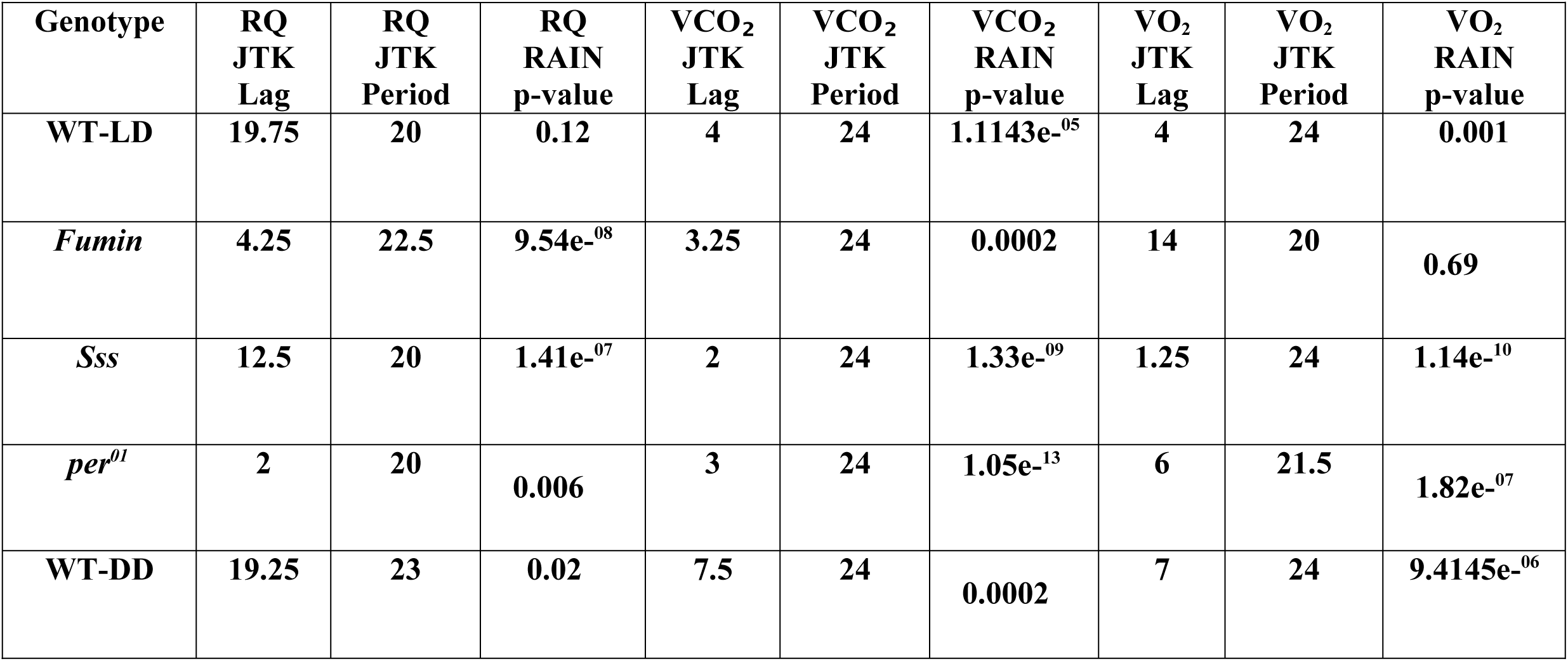
Rhythmicity metrics for respiratory parameters across genotypes.

| **Genotype** | **RQ**<br>**JTK**<br>**Lag** | **RQ**<br>**JTK**<br>**Period** | **RQ**<br>**RAIN**<br>**p-value** | **VCO<sub>2</sub>**<br>**JTK**<br>**Lag** | **VCO<sub>2</sub>**<br>**JTK**<br>**Period** | **VCO<sub>2</sub>**<br>**RAIN**<br>**p-value** | **VO<sub>2</sub>**<br>**JTK**<br>**Lag** | **VO<sub>2</sub>**<br>**JTK**<br>**Period** | **VO<sub>2</sub>**<br>**RAIN**<br>**p-value** |
| --- | --- | --- | --- | --- | --- | --- | --- | --- | --- |
| **WT-LD** | **19.75** | **20** | **0.12** | **4** | **24** | **1.1143e<sup>-05</sup>** | **4** | **24** | **0.001** |
| *Fumin* | **4.25** | **22.5** | **9.54e<sup>-08</sup>** | **3.25** | **24** | **0.0002** | **14** | **20** | **0.69** |
| *Sss* | **12.5** | **20** | **1.41e<sup>-07</sup>** | **2** | **24** | **1.33e<sup>-09</sup>** | **1.25** | **24** | **1.14e<sup>-10</sup>** |
| *per<sup>01</sup>* | **2** | **20** | **0.006** | **3** | **24** | **1.05e<sup>-13</sup>** | **6** | **21.5** | **1.82e<sup>-07</sup>** |
| **WT-DD** | **19.25** | **23** | **0.02** | **7.5** | **24** | **0.0002** | **7** | **24** | **9.4145e<sup>-06</sup>** |

### Diurnal Variation in CO₂ Production, O₂ Consumption, and Respiratory Quotient Across Genotypes

To investigate how sleep and diurnal regulation influence temporal patterns of metabolism, we measured carbon dioxide production (VCO₂), oxygen consumption (VO₂), and respiratory quotient (RQ) across day and night phases in wild-type flies maintained under light-dark cycles (WT-LD), wild-type flies kept in constant darkness (WT-DD) to assess free-running circadian regulation in the absence of external light-dark cues, short-sleep mutants (*fumin* [*fmn*] and *sleepless* [*sss*]), and the circadian clock mutant *period*^01^ (*per*^01^). The average values of VCO₂, VO₂ and RQ during the day (ZT0-12) and night (ZT12-24) are presented in **Figure S4**.

The *sss* mutant exhibited the highest metabolic rates among all genotypes, with significantly elevated VCO₂ and VO₂ during both day and night relative to WT-LD (p < 0.001). A pronounced day-night difference in *sss* further indicated a strong phase-dependent increase in respiratory activity (**Figure S4**). The *fmn* mutant also showed significantly elevated VCO₂ and VO₂ compared to WT-LD (p < 0.001), with a detectable but less marked diurnal variation than in *sss* (**Figure S4**). WT-LD flies exhibited a robust diurnal pattern, with significantly higher respiratory rates during the day (p < 0.001), consistent with LD-associated temporal regulation of energy expenditure (**Figure S4**).

The circadian mutant *per^01^*, lacking a functional molecular clock, displayed an attenuated and phase-altered respiratory pattern. Both VCO₂ and VO₂ were significantly reduced during the day (p < 0.05), with no significant changes at night, indicative of disrupted or misaligned metabolic patterning (**Figure S4**). WT-DD flies, maintained in constant darkness, exhibited the lowest overall respiratory activity. Despite the absence of environmental light cues, VCO₂ and VO₂ retained significant day-night differences (**Figure S4**), consistent with persistent free-running circadian control in constant darkness (with altered amplitude and/or phase relative to LD).

### Diurnal and Circadian Variations in CO₂ Production, O₂ Consumption, and Respiratory Quotient Across Genotypes

To evaluate genotype-dependent 24-h rhythmic regulation of metabolism, we assessed rhythmicity in carbon dioxide production (VCO₂), oxygen consumption (VO₂), and respiratory quotient (RQ) in wild-type flies under light-dark conditions (WT-LD), wild-type flies in constant darkness (WT-DD), short-sleep mutants (*fumin* [*fmn*] and *sleepless* [*sss*]), and the circadian clock mutant *period^01^* (*per^01^*). All mutant strains were assayed under LD. Time-course data were analyzed using JTK CYCLE and RAIN algorithms, with rhythmicity classified as statistically significant (p ≤ 0.05), trending (0.05 < p < 0.1), or not significant (p ≥ 0.1). Rhythmic phase was estimated using JTK lag values (**Figures 2-3**, **Table 1**).

VCO₂ rhythms were statistically significant in WT-LD, WT-DD, *fmn*, *sss* and *per^01^*(**Figure 2**, **Table 1**). These findings demonstrate that VCO₂ exhibits robust rhythmic oscillations under both light-dark and constant darkness conditions, and that rhythmicity persists even in the absence of a functional *period* gene. VCO₂ peak occurred near ZT ∼4 in WT-LD, ZT ∼7.5 in WT-DD, ZT ∼3.25 in *fmn*, ZT ∼3 in *per^01^*, and ZT ∼2 in *sss*, indicating genotype-specific shifts in respiratory phase (**Figure 3**, **Table 1**).

VO₂ rhythmicity was statistically significant in WT-LD, WT-DD, *sss* and *per^01^* while not significant in *fmn* (**Figure 2**, **Table 1**). These findings indicate that rhythmic regulation of oxygen consumption is evident in constant darkness and in the absence of a functional *period* gene but is reduced in a sleep mutant. VO₂ peaked near ZT ∼4 in WT-LD, ZT ∼7 in WT-DD, ZT ∼1.25 in *sss* and ZT ∼6 in *per^01^*, suggesting conserved phase alignment with all phases within ±3 h of WT-LD (**Figure 3**, **Table 1**).

RQ also showed genotype-dependent rhythmicity, with significant diurnal oscillations observed in *fmn, sss* and *per^01^*, while no significant rhythms were observed in WT-LD or WT-DD (**Figure 2**, **Table 1**). In the rhythmic genotypes, RQ peaked at ZT ∼4.25 in *fmn,* ZT ∼2 in *per^01^* and ZT ∼12.5 in *sss,* indicating distinct RQ phase alignment across the sleep mutants (spread ∼8 h apart), in contrast to the conserved phase alignment of respiratory output (VO₂) (**Figure 3**, **Table 1)**.

### Temporal Profiling of Respiratory Quotient in Wild-Type Flies Under Light-Dark Conditions

RQ values corresponding to each 2-hour Zeitgeber Time (ZT) point were extracted from the 5-minute-binned dataset. Building on this alignment, we explored temporal relationships by systematically shifting the RQ time series by −120, −60, −30, −15, −5, +5, +15, +30, +60, and +120 minutes relative to the metabolite dataset. The continuous respirometry time series showed an oscillatory day-night pattern in RQ; metabolomics was then used to relate time-matched and lagged metabolite dynamics to RQ patterns (**Figure 4**).

**Figure 4.**
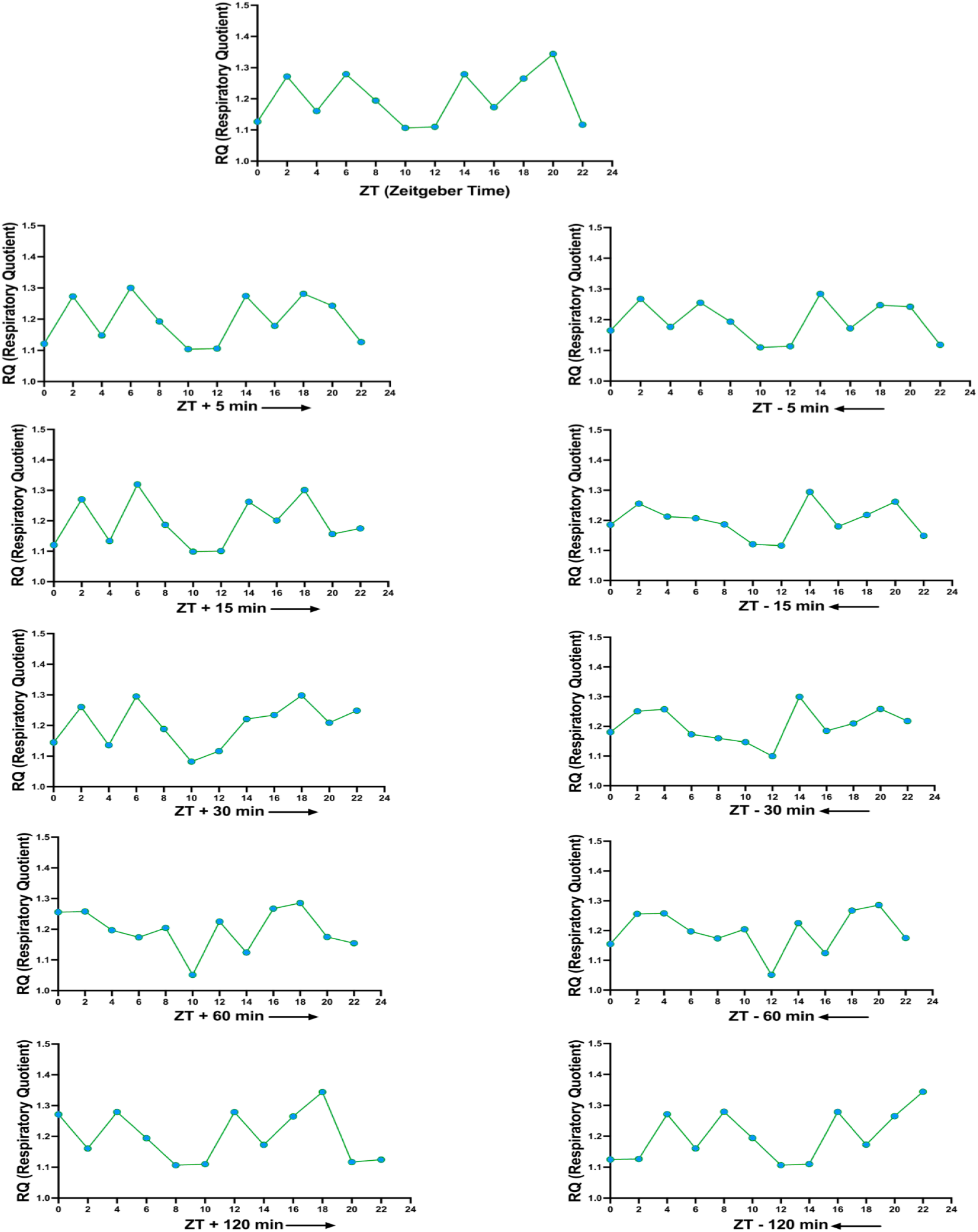
Temporal Profiling of Respiratory Quotient in Wild-Type Flies Under Light-Dark Conditions. Respiratory quotient (RQ) was extracted every 2 h across a 24-hour light-dark (LD) cycle in WT flies. Lag analyses were performed by advancing or delaying the RQ data by −120, −60, −30, −15, −5, +5, +15, +30, +60-, and +120-minutes relative to Zeitgeber Time (ZT). Each panel represents the RQ profile corresponding to a specific time shift, illustrating phase-dependent respiratory dynamics across the 24-hour LD cycle. Data represent approximately 300 flies per genotype across three experimental days (25 flies/chamber, four chambers/experiment). The chamber was used as the experimental unit; *n* denotes the number of chambers. RQ rhythmicity itself was assessed from the continuous respirometry time series (Table 1); Figure 4 shows the day-night RQ pattern used as the reference for the lag-based metabolite correlations, to which the metabolomics data contribute.

### Correlation of Respiratory Quotient with Metabolite Profiles and Temporal Lag Analysis in WT-LD Flies

To investigate the association between respiratory quotient (RQ) and metabolite in WT-LD flies, we initially performed Spearman correlation analysis (ρ) across a range of temporal lags (−120 to +120 minutes). Several metabolites demonstrated strong correlations with RQ (|ρ| > 0.7, *p* < 0.05), suggesting tight coupling between metabolic fluctuations and respiratory output. The full set of metabolites meeting this threshold across WT-LD, *fmn*, *sss*, *per^01^*, and WT-DD conditions, along with best lag direction, lag time, and Spearman ρ value and BH FDR, is provided in **Supplementary Table 1**. To minimize the influence of outliers and enhance data visualization, selected correlation patterns were subsequently presented as rank-based scatter plots. Representative examples are shown in **Figure 5a** to illustrate the metabolite-RQ correlation and lag-analysis workflow. Hydroxyhexadecenoylcarnitine and quinolinate are among the strongest positively- and negatively lagged RQ-correlated metabolites in WT-LD (ρ = +0.78 at +120 min lag and ρ = −0.77 at −120 min lag; **Supplementary Table 1**), illustrating the two opposite lag directions of the workflow. The full set of metabolites showing strong RQ-associated correlations across WT-LD, *fmn*, *sss*, *per^01^*, and WT-DD conditions is provided in **Supplementary Table 1**. We define ‘anticipatory’ as metabolite changes that precede the associated respiratory shift (negative lag) and ‘reactive’ as changes that follow or coincide with the respiratory shift (positive lag), and we use these terms consistently throughout. To further characterize these associations, metabolites were categorized based on the timing of their peak changes relative to RQ fluctuations, exhibiting either positive or negative lag. To enable direct comparison of temporal dynamics between RQ and metabolite profiles across ZT 0 to ZT 24, z-scoring was performed to normalize differences in absolute magnitude and measurement units. Within this framework, a positive lag indicates that metabolite changes occur after shifts in RQ, requiring the metabolite profile to be shifted forward (rightward) along the time axis for optimal alignment. Conversely, a negative lag signifies that metabolite changes precede RQ fluctuations, necessitating a backward (leftward) shift of the metabolite profile to achieve alignment (**Figure 5b**).

**Figure 5.**
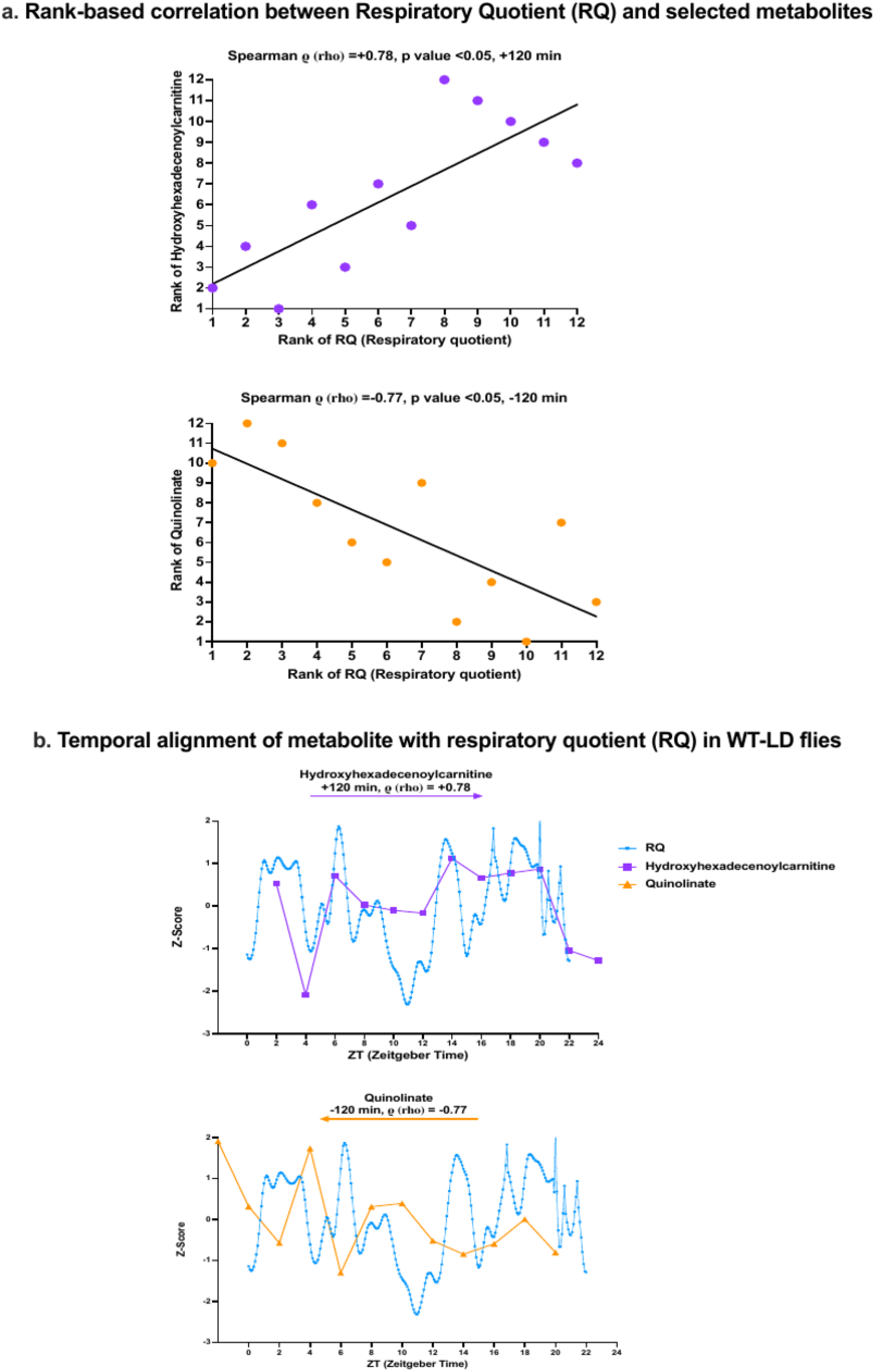
***a. Rank-based correlation between Respiratory Quotient (RQ) and selected metabolites.*** Scatter plots depict the rank-order relationships between RQ and (A) Hydroxyhexadecenoylcarnitine and (B) Quinolinate. Each point represents a timepoint after aligning data with respective time shifts. Spearman’s rank correlation coefficient (ρ) and corresponding p-values are indicated on each panel. A positive lag (+120 minutes for Hydroxyhexadecenoylcarnitine) or negative lag (−120 minutes for Quinolinate) denotes the temporal shift in RQ relative to the metabolite dataset. Statistical significance was defined as p < 0.05. ***b. Temporal alignment of metabolite with respiratory quotient (RQ) in WT-LD flies.*** Z-scored time series of RQ, Hydroxyhexadecenoylcarnitine, and Quinolinate across a 24-hour light-dark cycle. **Top Panel:** Hydroxyhexadecenoylcarnitine shows a positive lag of +120 minutes and strong positive correlation with RQ (ρ=+0.78), suggesting its rise after changes in RQ. **Bottom Panel:** Quinolinate shows a negative lag of –120 minutes and strong negative correlation with RQ (ρ=-0.77), indicating it precedes changes in RQ.

### Metabolite Response Dynamics Relative to RQ and Functional Pathway Enrichment of RQ-Associated Metabolites in WT-LD Flies

To further explore the temporal relationship between the metabolites and respiratory quotient (RQ), we generated a clustered heatmap of metabolites significantly correlated with RQ. Metabolites were filtered based on statistical significance (|ρ| > 0.7, p < 0.05 at any timepoint), and their temporal profiles were realigned relative to respiration, anchoring RQ at t = 0 ( **Figure 6, Supplementary Table 2**). This flipping step was essential to standardize the visualization of metabolites either preceding or following respiratory changes, facilitating clearer interpretation of biological timing relationships.

**Figure 6.**
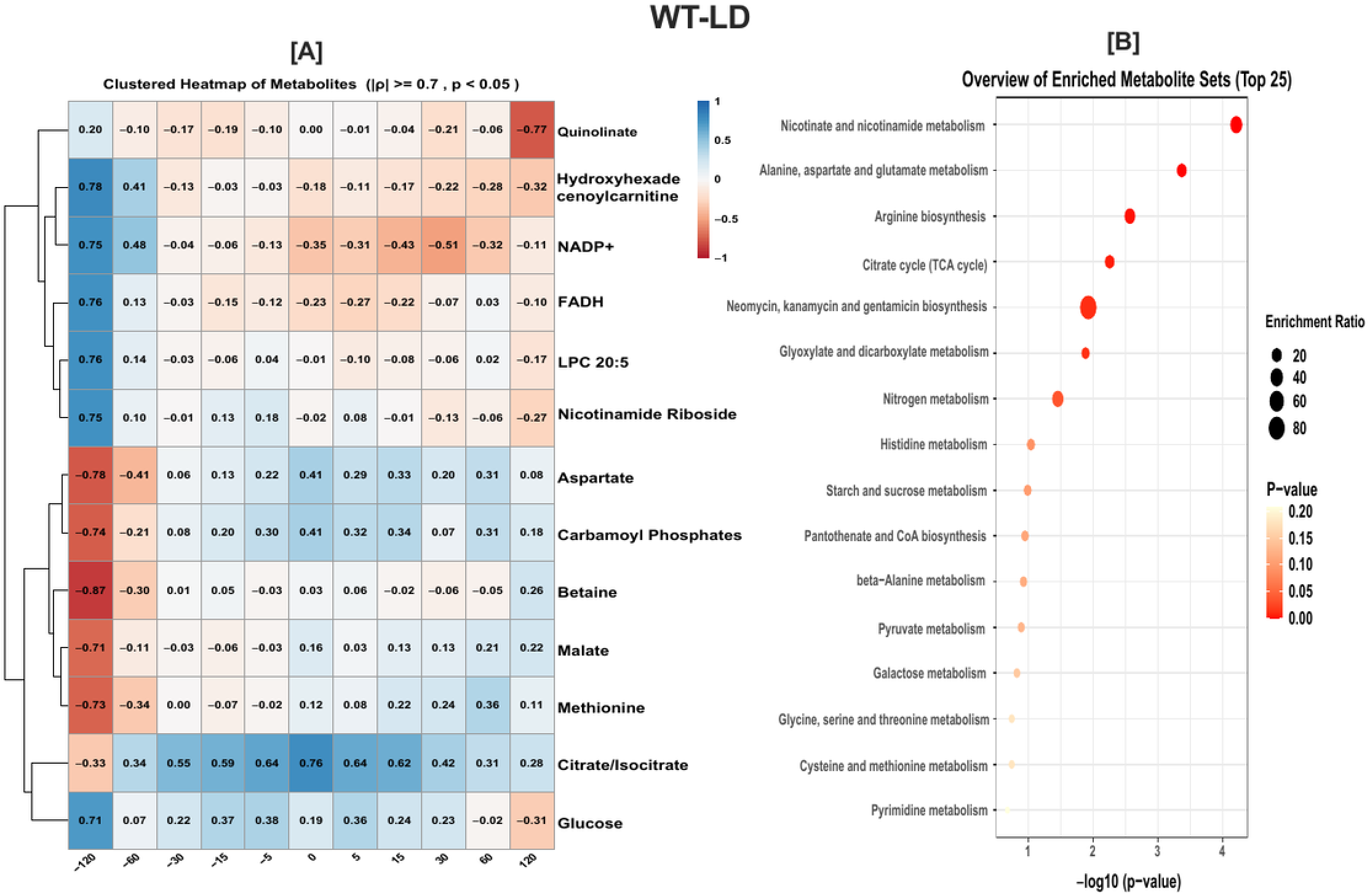
Metabolite-Respiratory Quotient Correlations and Pathway Enrichment in WT-LD Flies. (A) Heatmap depicting Spearman correlations (|ñ| > 0.7, p < 0.05) between respiratory quotient (RQ) and metabolite across the 24-hour LD cycle in wild-type flies maintained under light-dark (LD) conditions. Metabolites were clustered based on correlation patterns, highlighting groups with similar temporal associations with respiratory activity. (B) Pathway enrichment analysis of significantly correlated metabolites, illustrating metabolic pathways most closely linked to respiratory dynamics. Pathways with a p-value less than 0.05 were considered statistically significant.

To identify key metabolic pathways associated with diurnal regulation of respiration, we next performed pathway enrichment analysis on the RQ-correlated metabolites using MetaboAnalyst. Enriched pathways were identified at a significance threshold of p < 0.05. WT-LD-specific pathway enrichment results are presented in **Figure 6**. Consistent with these findings, WT flies under LD conditions exhibited diurnal-regulated metabolic rhythms, characterized by coordinated activity in amino acid metabolism, redox balance, and energy-generating pathways such as the TCA cycle and glyoxylate metabolism.

### RQ-Linked Metabolic Shifts and Pathway Enrichment in Sleep Mutants

We applied the same analysis method to sleep mutant flies (*fmn* and *sss*) to identify metabolic pathways associated with altered respiratory rhythms. Metabolites significantly correlated with RQ (|ρ| > 0.7, *p* < 0.05) were identified, and their temporal profiles were realigned relative to respiration by anchoring RQ at *t* = 0. Clustered heatmaps were generated to visualize metabolite response dynamics, and pathway enrichment analysis of RQ-associated metabolites was conducted using MetaboAnalyst (*p* < 0.05) (**Figure 7**). This analysis revealed that sleep mutants exhibit genotype-specific metabolic disruptions implicating mitochondrial and energy-related pathways. In *fmn* flies, alterations were observed in arginine biosynthesis, purine and nitrogen metabolism, and butanoate pathways, suggestive of dysregulated nitrogen balance and mitochondrial dysfunction. In contrast, *sss* mutants showed perturbations in glycine, serine, and threonine metabolism, as well as carbohydrate and antibiotic biosynthetic pathways, reflecting altered carbon utilization and mitochondrial-associated metabolic stress.

**Figure 7.**
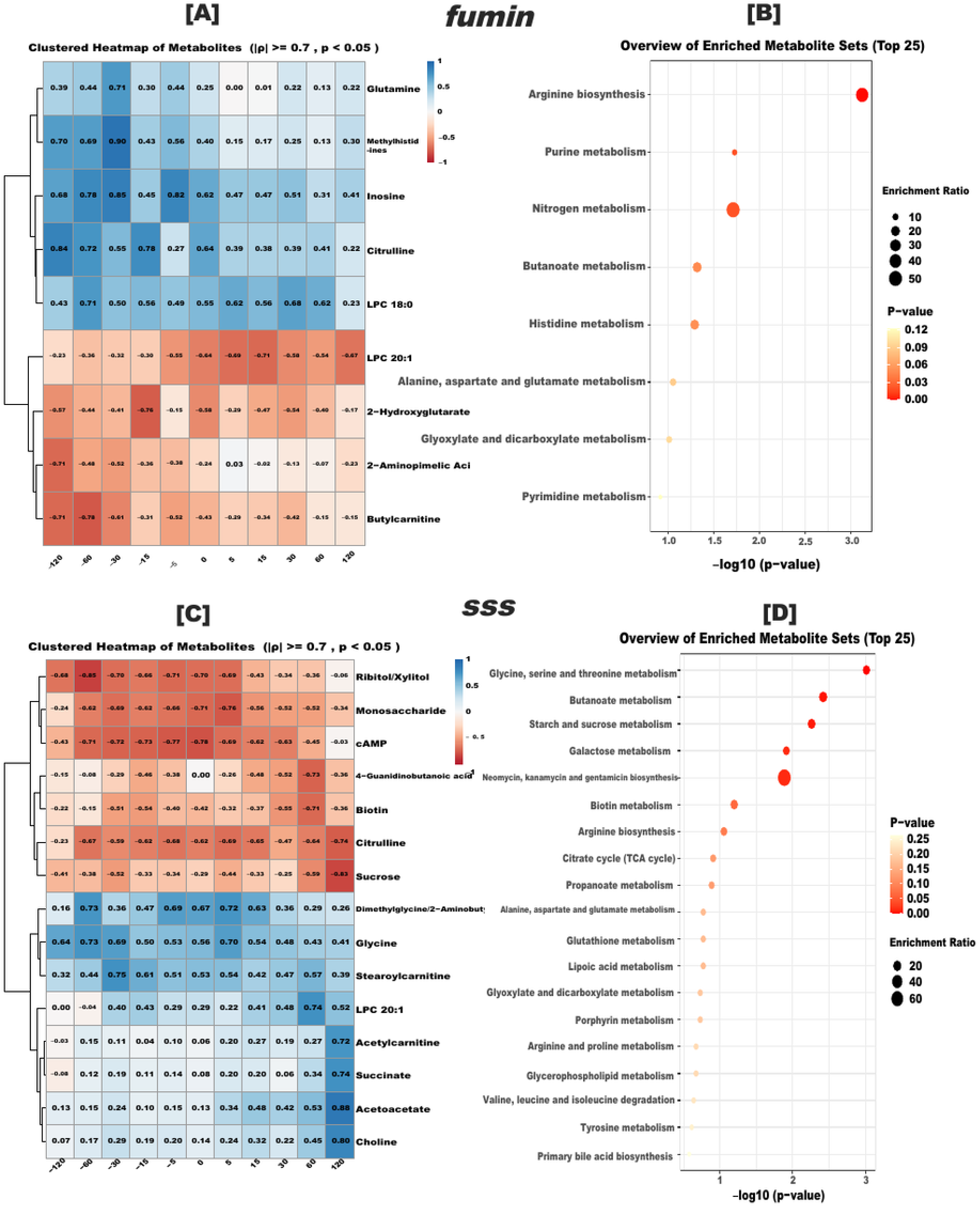
Correlation of Metabolites with Respiratory Quotient and Pathway Enrichment in Short-Sleep Mutants. (A, C) Heatmaps showing Spearman correlations (|ρ| > 0.7, p < 0.05) between respiratory quotient (RQ) and metabolites in the short-sleep mutants *fmn* (A) and *sss* (C). (B, D) Pathway enrichment analyses of metabolites significantly correlated with RQ in *fmn* (B) and *sss* (D). Pathways with p-values less than 0.05 were considered statistically significant.

### RQ-Linked Metabolic Shifts in Circadian Mutants and WT Flies in Constant Darkness

Circadian clock mutant flies (*per^01^*) and wild-type flies maintained under constant darkness (WT-DD) were analyzed using the same approach. We used constant darkness (DD) to assess free-running circadian regulation in the absence of external light–dark cues. Metabolites exhibiting strong correlations with RQ (|ρ| > 0.7, p < 0.05) were identified and temporally aligned by anchoring RQ at t = 0. Clustered heatmaps were constructed to visualize the dynamic responses of these metabolites, and pathway enrichment analysis of RQ-associated metabolites was performed using MetaboAnalyst (p < 0.05) (**Figure 8**). Both *per^01^* and WT-DD flies exhibited disrupted temporal coordination of mitochondrial metabolism. *per^01^* mutants showed dysregulation in amino acid metabolism, purine turnover, and redox-associated pathways such as glutathione and nicotinate metabolism, consistent with impaired redox buffering and energy imbalance. In contrast, WT-DD flies exhibited alterations in sulfur- and nitrogen-containing amino acid pathways, indicative of desynchronized but intact metabolic cycling in the absence of external light cues.

**Figure 8.**
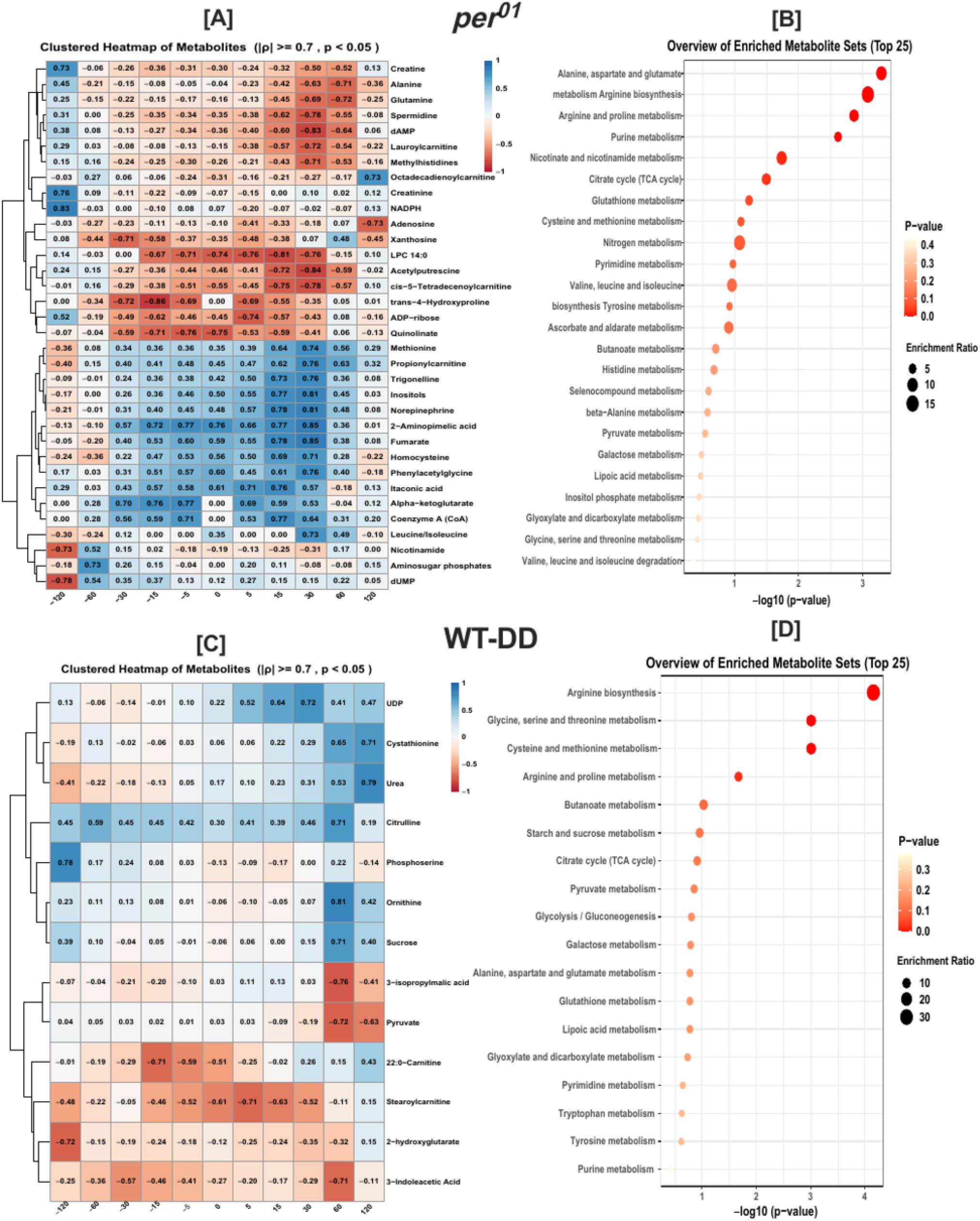
Correlation of Metabolites with Respiratory Quotient and Pathway Enrichment in Circadian Mutant and Wild-Type Flies in Constant Darkness. (A, C) Heatmaps displaying Spearman correlations (|ρ| > 0.7, p < 0.05) between respiratory quotient (RQ) and metabolites in the circadian mutant *per^01^* (A) and wild-type flies maintained in constant darkness (WT-DD) (C). (B, D) Pathway enrichment analyses of metabolites significantly correlated with RQ in *per^01^* (B) and WT-DD (D). Pathways with p-values less than 0.05 were considered statistically significant.

### Gut Tissue Respirometry Reveals Elevated Baseline Mitochondrial Respiration in fmn and per^01^ Mutants

Because pathway-level metabolomics implicated mitochondrial pathways in sleep and circadian mutants, we directly assayed mitochondrial respiration by measuring baseline oxygen consumption rate (OCR) in dissected gut tissue from *fmn* and *per^01^* flies relative to iso controls. Both *fmn* and *per^01^* guts showed significantly elevated baseline OCR (*fmn* vs *iso^31^*, n = 12 vs 12; *per^01^*vs *iso^31^*, n = 12 vs 11 across 3 runs; Mann-Whitney test, p < 0.05; Figure 9A,B). Because only baseline OCR was measured, we interpret this as altered (elevated) baseline mitochondrial respiration rather than reduced capacity or a specific coupling defect (**Figure 9**).

**Figure 9.**
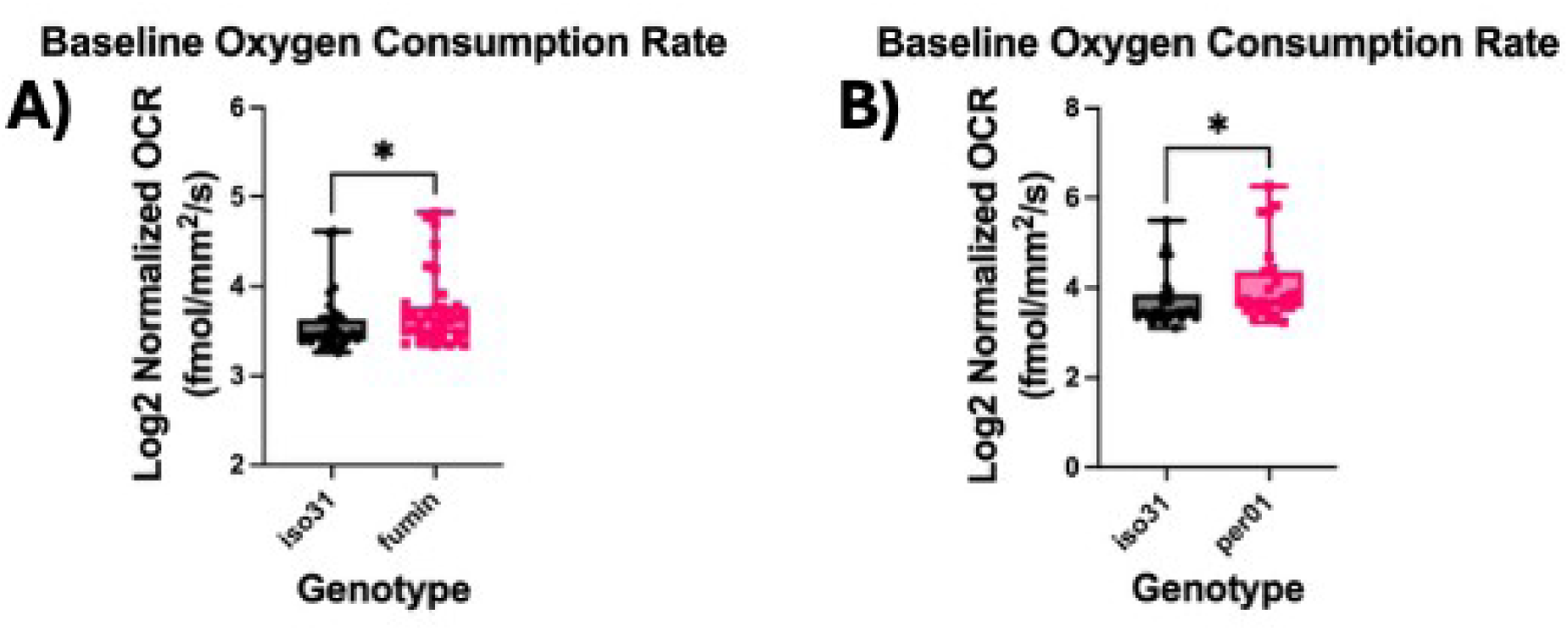
Baseline oxygen consumption in isolated gut tissue from fmn and per^01^ mutants. Baseline oxygen consumption rate (OCR, fmol/mm²/s) measured by tissue respirometry in individual dissected guts from the short-sleep mutant *fmn* (A) and the circadian-clock mutant *per^01^* (B) relative to *iso^31^* genetic-background controls. For each gut, OCR was averaged over a 24-hour window beginning ∼12 h after loading, once tissue respiration had stabilized (12–36 h). To combine independent runs, OCR was normalized to the median of the *iso^31^* controls within each run and log2(x+10)-transformed. One *iso^31^* gut in the third run of the *per^01^* experiment was excluded as an outlier; no other values were removed. Baseline OCR differed significantly between *iso^31^* and each mutant (Mann-Whitney test, p < 0.05; pooled across 3 independent runs; *fmn* vs *iso^31^*, n = 12 vs 12; *per^01^* vs *iso^31^*, n = 12 vs 11 after the *iso^31^* outlier exclusion).

## DISCUSSION

### Temporal Misalignment Alters Fuel Utilization and Respiratory Rhythms

Our integrative approach, combining whole-organism respirometry [18, 26] with LC-MS-based metabolomics [27, 36-40], reveals how sleep and circadian disruptions distinctly alter energy metabolism in *Drosophila melanogaster*. By comparing wild-type flies with mutants affecting sleep (*fumin* and *sleepless*) [21, 22, 41, 42] and circadian clock function (*per^01^*) [24], we have captured genotype-specific differences in respiratory dynamics and substrate usage across daily cycles under tightly controlled environmental conditions. These findings establish a functional framework for understanding how behavioral state and clock integrity shape metabolic homeostasis.

Across the manuscript, the central takeaway is that wild-type flies under LD exhibit anticipatory alignment of fuel selection with time of day, whereas short-sleep mutants (*fmn, sss*) and clock-disrupted flies (*per^01^*) show reactive or misaligned metabolism under our conditions. We therefore focus the Discussion on loss of temporal coordination between respiratory output and pathway-level metabolism, rather than reiterating rate changes alone.

Metabolites in wild-type LD typically showed peak correlations preceding respiratory changes (negative lag), reflecting anticipatory diurnal metabolic regulation under LD conditions. In contrast, mutants displayed a shift to reactive correlations (positive lag), indicating loss of temporal synchronization and proactive metabolic adjustments. These phenotypes parallel mammalian systems, where sleep loss and circadian misalignment are linked to elevated basal metabolic rate, a shift toward carbohydrate oxidation and lipid/protein catabolism, and blunted, phase-shifted respiratory oscillations [3, 43, 44]. In our flies, the short-sleep mutants show the analogous catabolic shift (*fmn* RQ 1.09; *sss* RQ 0.94), and *per^01^* and WT-DD show dampened, phase-shifted respiratory rhythms [45, 46]. These conserved responses underscore the translational value of *Drosophila* for dissecting the interplay between sleep, circadian timing, and metabolic regulation, while physiological differences between flies and mammals caution against direct mechanistic extrapolation.

### Wild-Type Flies Exhibit Diurnal Regulation of Biosynthesis and Redox Balance Under LD Cycles

In wild-type flies maintained under a 12:12 light-dark (LD) cycle, RQ-correlated metabolites revealed tightly coordinated diurnal regulation of biosynthetic and redox pathways. An increase in nicotinate and nicotinamide metabolism-highlighted by changes in quinolinate, nicotinamide riboside, and L-aspartic acid-suggests a direct link between mitochondrial respiration and NAD⁺ biosynthesis [47, 48]. These NAD⁺ precursor changes occur before respiration increases (negative lag), highlighting anticipatory NAD⁺ regulation. Concurrent alterations in alanine, aspartate, and glutamate metabolism (via citric acid and carbamoyl phosphate) and arginine biosynthesis indicate dynamic alignment of amino acid turnover with energy demand and nitrogen disposal [49, 50]. RQ-associated enrichment of citrate cycle intermediates such as malate and citric acid further supports diurnal gating of oxidative phosphorylation [51]. Elevated levels of D-glucose were mapped to the KEGG pathway for neomycin, kanamycin, and gentamicin biosynthesis. While *Drosophila* does not synthesize these antibiotics, the enrichment likely reflects the presence of shared carbohydrate intermediates (e.g., glucose, glucose-6-phosphate, UDP-glucose) common to multiple biosynthetic processes. This mapping is more indicative of altered carbohydrate and amino sugar metabolism, or potentially reflects microbial contributions, rather than actual antibiotic production by the host [52, 53]. Additional contributions from glyoxylate and dicarboxylate metabolism reinforce the role of anaplerotic flux and redox cycling [54]. Notably, persistent correlations between RQ and pantothenate precursors suggest synchronized CoA biosynthesis, potentially coordinating fatty acid oxidation and TCA cycle input [55]. Together, these results demonstrate that under LD conditions, wild-type flies sustain temporal coordination between respiration and mitochondrial metabolism, integrating redox homeostasis, nitrogen metabolism, and substrate availability in an LD-associated diurnal manner.

### Altered Amino Acid Metabolism and Mitochondrial Dysregulation in fmn Mutants

The *fumin* (*fmn*) mutant, characterized by chronic sleep loss, exhibited genotype-specific disruptions in metabolic coordination, as revealed by metabolites whose temporal dynamics were strongly correlated with respiratory quotient (RQ). In *fmn* flies, significant RQ correlations (*p* < 0.05) were identified in pathways including arginine biosynthesis (citrulline, glutamine), purine metabolism (inosine, glutamine), and nitrogen metabolism-suggesting a decoupling of amino acid turnover and nucleotide cycling from respiratory demand [56, 57]. Glutamine’s strong association with RQ highlights its central role in buffering nitrogen flux and supporting biosynthetic processes under energetic stress [58]. Correlated dynamics of 2-hydroxyglutarate, a key intermediate in butanoate metabolism, point to impaired TCA cycle flux and redox imbalance [59]. Likewise, RQ-associated shifts in methylhistidine suggest altered histidine metabolism, potentially impacting mitochondrial protein turnover and methylation processes. Acylcarnitine, indicators of lipid oxidation, showed consistently strong RQ correlations across genotypes, most pronounced in *fmn*, underscoring lipid metabolism’s role in respiratory coupling. Particularly in mutants, increased reliance on fatty acid oxidation may reflect compensatory responses to disrupted carbohydrate metabolism and sustained energetic demands. Together, these findings indicate a failure in aligning substrate oxidation with mitochondrial respiratory output in *fmn* mutants. Building on prior evidence of mitochondrial stress in sleep-deprived *fmn* flies [60], our integrative respirometry-metabolomics analysis reveals disrupted coupling between energy metabolism and amino acid pathways, underscoring the physiological cost of chronic sleep loss on mitochondrial homeostasis. These interpretations are further supported by gut-tissue respirometry showing elevated baseline mitochondrial respiration (OCR) in *fmn* relative to iso31 controls. Together with prior evidence of ROS accumulation (oxidative stress) in *fmn* [60], these functional data indicate that chronic sleep loss in *fmn* is associated with altered mitochondrial respiration.

### Disrupted Carbon and Amino Acid Metabolism in sss Mutants

The *sleepless* (*sss*) mutant displayed significant disruptions in carbon and amino acid metabolism, as indicated by strong correlations between respiratory quotient (RQ) and metabolites involved in glycine, serine, and threonine metabolism (choline, dimethylglycine, glycine), butanoate metabolism (acetoacetate, succinate), and carbohydrate pathways [22]. Notably, RQ-associated shifts in choline and dimethylglycine suggest dysregulation of one-carbon metabolism and methyl group transfer, which can impair mitochondrial function and epigenetic stability [61, 62]. Correlated dynamics of acetoacetate, a ketone body often found in starvation, and succinate (both key intermediates of butanoate metabolism) are consistent with altered TCA cycle input and redox balance [63, 64]. In parallel, strong RQ correlations with sucrose and D-glucose (from both starch/sucrose and galactose metabolism) indicate disrupted carbohydrate processing and inefficient substrate utilization [65]. cAMP and biotin were also found to associate with respiration, both unique in metabolic regulation through altered neuromodulatory signaling and cofactor metabolism. These findings suggest that *sss* mutants exhibit altered metabolic routing in response to respiratory demand, shifting substrate utilization toward carbohydrate and amino acid pathways under sleep-deprived conditions, consistent with metabolic stress responses observed in other models of sleep loss [66].

### Loss of Temporal Coupling of Mitochondrial Metabolism in Clock Mutants

In *period* (*per^01^*) mutants lacking a functional circadian clock, the temporal coordination between metabolism and respiration was disrupted across nearly every measured metabolite through intermittent strong correlations with respiration, reflecting broad metabolic chaos [1, 67]. Strong correlations with RQ were observed in alanine, aspartate, and glutamate metabolism (L-alanine, glutamine, fumarate, oxoglutarate) and arginine biosynthesis, suggesting inefficient routing of amino acid-derived substrates into mitochondrial energy production [50]. Delayed RQ-associated dynamics of glutamine, oxoglutarate, and fumarate indicate a mismatch between substrate availability and respiratory output [2]. Additionally, metabolites from arginine and proline metabolism (creatine, spermidine, N-acetylputrescine, 4-hydroxyproline) exhibited altered RQ correlations, suggesting impaired mitochondrial redox buffering and compromised integrity [68]. Perturbations in purine metabolism-including adenosine, deoxyadenosine monophosphate, xanthosine, and ADP-ribose-further point to dysregulated ATP turnover and nucleotide imbalance. Correlations with quinolinic acid and niacinamide reflect altered nicotinate and nicotinamide metabolism, implicating disrupted NAD⁺ biosynthesis and redox state [48]. Additional RQ-associated changes in NADPH and TCA intermediates (fumarate, oxoglutarate) align with impaired glutathione metabolism and altered mitochondrial metabolism. Gut-tissue respirometry in *per^01^* showed elevated baseline OCR. Because only baseline respiration was measured, we interpret this as altered (elevated) baseline mitochondrial respiration — which can reflect inefficient or uncoupled respiration, or compensatory upregulation, rather than increased functional capacity — consistent with the metabolomic signatures of disrupted redox balance and mitochondrial metabolism. Microbial-derived metabolites (trigonelline, phenylacetylglycine) were also identified in the correlation analysis, suggesting potential gut microbiome interactions that become evident in circadian disruption. Together, these findings suggest that the loss of circadian timing in *per^01^* mutants lead to widespread uncoupling of substrate utilization and mitochondrial respiration. This breakdown in temporal regulation disrupts energy homeostasis, highlighting the essential role of the circadian clock in coordinating metabolic flux with respiratory demand [1, 51, 69].

### Metabolic Desynchrony and Redox Imbalance in Wild-Type Flies Under Constant Darkness

In wild-type flies maintained under constant darkness (DD), the absence of environmental light cues led to disrupted temporal alignment between mitochondrial respiration and metabolic pathways [70, 71]. Generally, we note a shift towards reactive metabolic adjustments (positive lags) compared to WT-LD instead of anticipatory preparation, indicating reduced metabolic efficiency. RQ-correlated metabolites showed significant enrichment in arginine biosynthesis (citrulline, ornithine, urea), glycine, serine, and threonine metabolism (L-cystathionine, phosphoserine, pyruvic acid), and cysteine and methionine metabolism-indicating altered integration of nitrogen and sulfur amino acid metabolism with respiratory activity [49]. Notably, the convergence of pyruvic acid across multiple pathways suggests dysregulated entry points into the TCA cycle, while the consistent association of L-cystathionine highlights impaired redox buffering and methylation capacity [72, 73]. Altered correlations in arginine and proline metabolism (ornithine, pyruvic acid) further underscore compromised coupling between amino acid turnover and mitochondrial energy production [29]. Altered lipid metabolism and increased nitrogen catabolism are likely compensatory mechanisms in the absence of environmental cues. These findings build on previous reports of circadian desynchrony by providing functional evidence that environmental light cues contribute to coherent coordination between respiration and key metabolic circuits [50, 74]. The resulting metabolic misalignment under constant darkness emphasizes the circadian clock’s dependence on external entrainment to regulate energy homeostasis at the organismal level [54]. Our DD data show that endogenous free-running circadian regulation persists but that removing external light–dark cues degrade the temporal coordination between respiration and metabolism, indicating that external entrainment normally strengthens this coordination under our conditions; however, we acknowledge that the coupling may be different in mammals. We also restrict our conclusions to the genotypes and conditions tested (*fmn, sss, and per^01^*) and do not generalize these effects to acute sleep deprivation, which was not performed in this study. Accordingly, the ‘metabolic misalignment’ observed in constant darkness (DD) likely results from a decrease in synchrony due to the removal of external light:dark cues. We also acknowledge the limits of this replication: with n = 12 chambers per genotype, statistical power to detect small-magnitude differences and subtle phase shifts is limited, and negative calls (e.g., arrhythmicity) should be interpreted with this caveat.

Locomotor activity and feeding could not be measured during the respirometry recordings, so genotype differences in respiratory parameters should be interpreted with caution. To assess whether behavioral timing could account for these differences, we compared our respiratory rhythms with the feeding and activity rhythms reported for these genotypes in our previous work [75]: wild-type feeding peaked at ZT ∼3.25 and *fmn* feeding was phase-delayed to ZT ∼4.5, while *fmn* also showed elevated locomotor activity, particularly during the dark period. In our data the *fmn* VCO_2_ rhythm peaks at ZT ∼3.25 (RQ at ZT ∼4.25), so the respiratory peak slightly precedes the feeding peak; the phase of the *fmn* respiratory rhythm is therefore not driven by feeding, although the elevated activity of *fmn* may contribute to its higher overall metabolic rate. Feeding and activity were not measured for *sss*.

Because only males were analyzed, our conclusions may not generalize to females, which can show sex-specific metabolic physiology. Inclusion of female flies and direct sex comparisons across LD and DD conditions will be an important future direction. More specifically, females carry a higher reproductive and biosynthetic load (egg production) that typically raises metabolic rate and can shift RQ toward lipogenesis and alter the amplitude and phase of diurnal respiratory rhythms; females might therefore show larger or differently timed effects than the males studied here.

## CONCLUSIONS

This study demonstrates that sleep loss and circadian disruption impair metabolic homeostasis in *Drosophila melanogaster* via distinct yet converging mechanisms. Wild-type flies under light-dark cycles (WT-LD) maintained coordinated anticipatory respiratory and metabolic rhythms, while sleep mutants (*fmn*, *sss*) and circadian-disrupted flies (*per^01^*) showed altered substrate utilization, redox imbalance, and uncoupling of mitochondrial pathways, likely reactive to respiration. These findings highlight the essential role of both sleep and circadian timing in regulating energy metabolism. The presence of neuromodulatory metabolites (e.g., cAMP in *sleepless* mutants) and microbial-derived compounds (trigonelline, phenylacetylglycine in *per^01^* mutants) points toward additional layers of metabolic regulation influenced by neuronal activity and gut microbiota interactions, revealing more complex metabolic dysregulation in sleep-deprived and circadian-disrupted states. By combining high-resolution respirometry with targeted metabolomics, we establish *Drosophila* as a powerful model for investigating the molecular basis of sleep- and clock-related metabolic dysfunction and potential therapeutic interventions.

## Author Contributions

**Conceptualization, F.A., A.S. and A.M.W.; methodology, F.A., D.M.M., P.H., J.K. A.N. and A.M.W.; formal analysis, F.A., A.S.G., A.N. and A.M.W.; investigation, F.A. A.N. and D.M.M.; resources, A.S. and A.M.W.; writing-original F.A.; writing-review and editing, F.A., A.S.G., J.K., A.S., and A.M.W.; visualization, F.A., A.S.G, and A.M.W.; supervision, A.M.W.; and funding acquisition, A.S. and A.M.W.**

## Notes

The authors declare no competing financial interest.

## ACKNOWLEDGMENTS

We thank Pinky Kain for assistance with *Drosophila* stock maintenance and insightful discussions on *Drosophila* mutants. We are also grateful to Sara B. Noya for her support during respirometry calibration.

## SUPPLEMENTARY MATERIALS

### Supplementary Figure Legends and Tables

**Figure S1.**
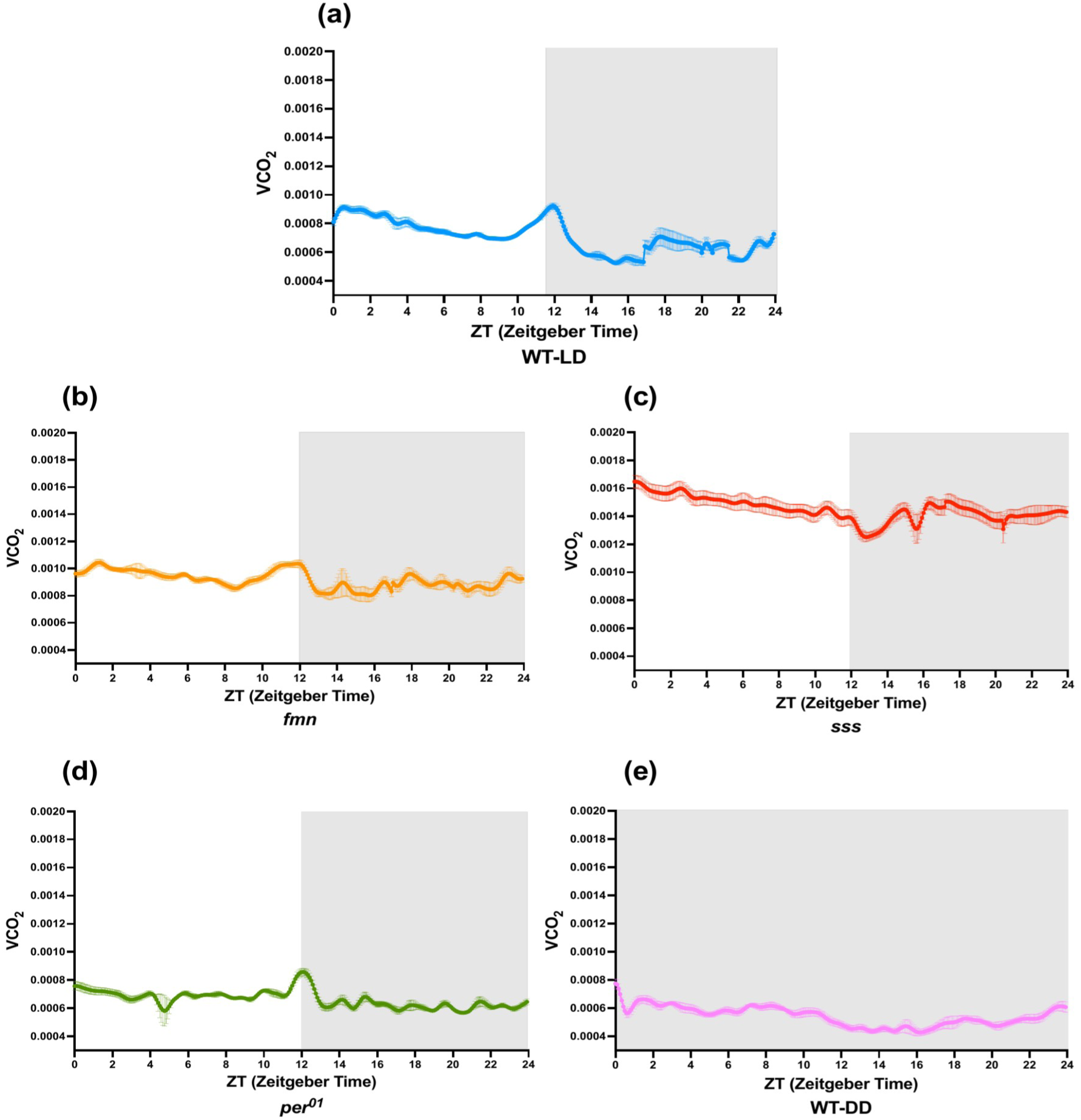
a-e. VCO₂ profiles across the 24-hour recording period in different genotypes. VCO₂ was continuously measured at one-second intervals from Zeitgeber Time (ZT) 0 to 24 and averaged into 5-minute bins for analysis. The traces display mean VCO₂ values across the 24-hour recording period for the following genotypes: wild-type flies under light-dark conditions (WT-LD), short-sleep mutants *fumin* (*fmn*) and *sleepless* (*sss*), the circadian clock mutant *period*^01^ (*per^01^*), and wild-type flies in constant darkness (WT-DD). For all panels, traces are presented as mean ± SEM.. Data represent approximately 300 flies per genotype across three experimental days (25 flies/chamber, four chambers/experiment). The chamber was used as the experimental unit; *n* denotes the number of chambers.

**Figure S2.**
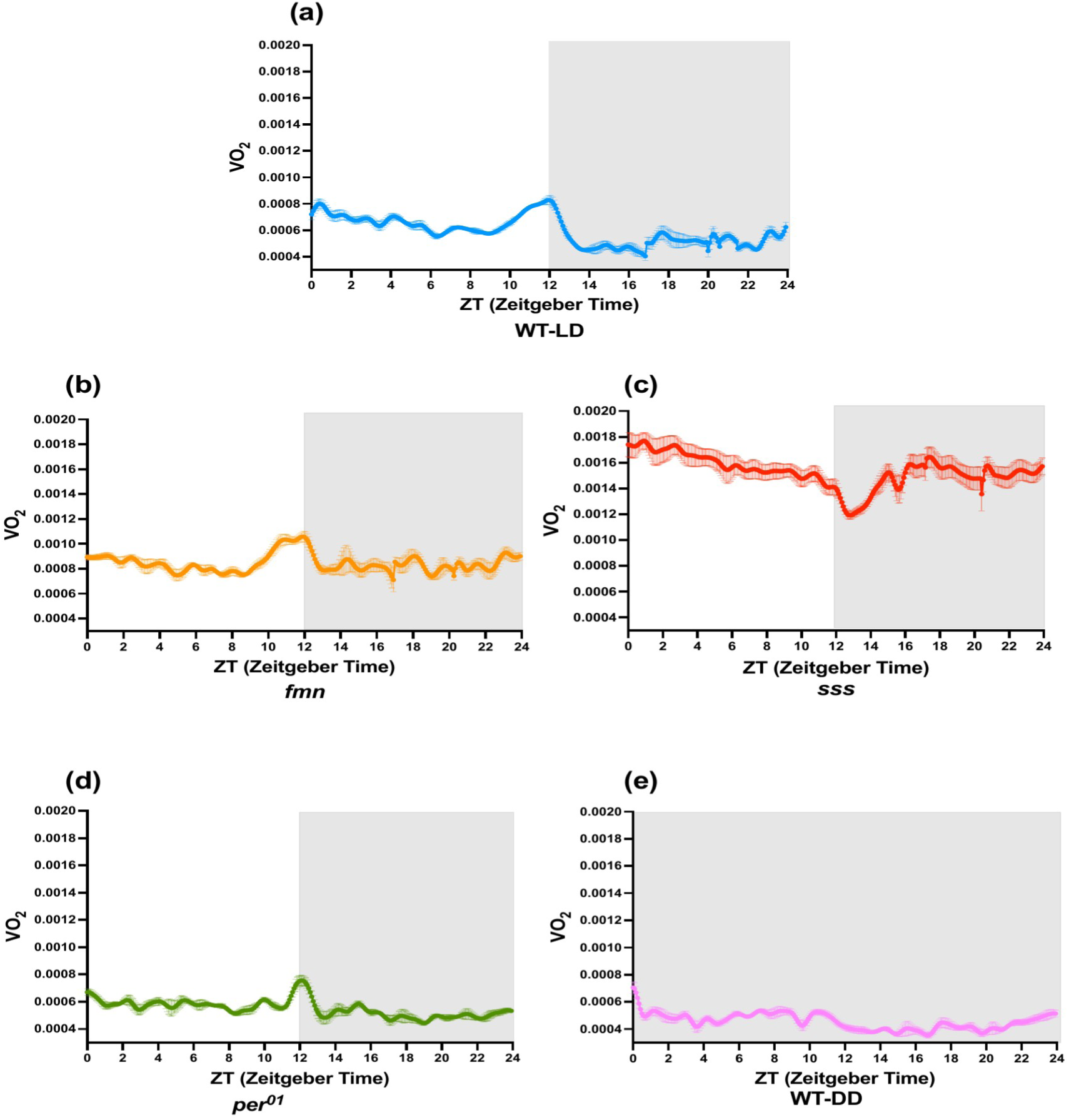
a-e. VO₂ profiles across the 24-hour recording period in different genotypes. VO₂ was continuously measured at one-second intervals from Zeitgeber Time (ZT) 0 to 24 and averaged into 5-minute bins for analysis. The traces display mean VO₂ values across the 24-hour recording period for the following genotypes: wild-type flies under light-dark conditions (WT-LD), short-sleep mutants *fumin* (*fmn*) and *sleepless* (*sss*), the circadian clock mutant *period*^01^ (*per^01^*), and wild-type flies in constant darkness (WT-DD). For all panels, traces are presented as mean ± SEM. Data represent approximately 300 flies per genotype across three experimental days (25 flies/chamber, four chambers/experiment). The chamber was used as the experimental unit; *n* denotes the number of chambers.

**Figure S3.**
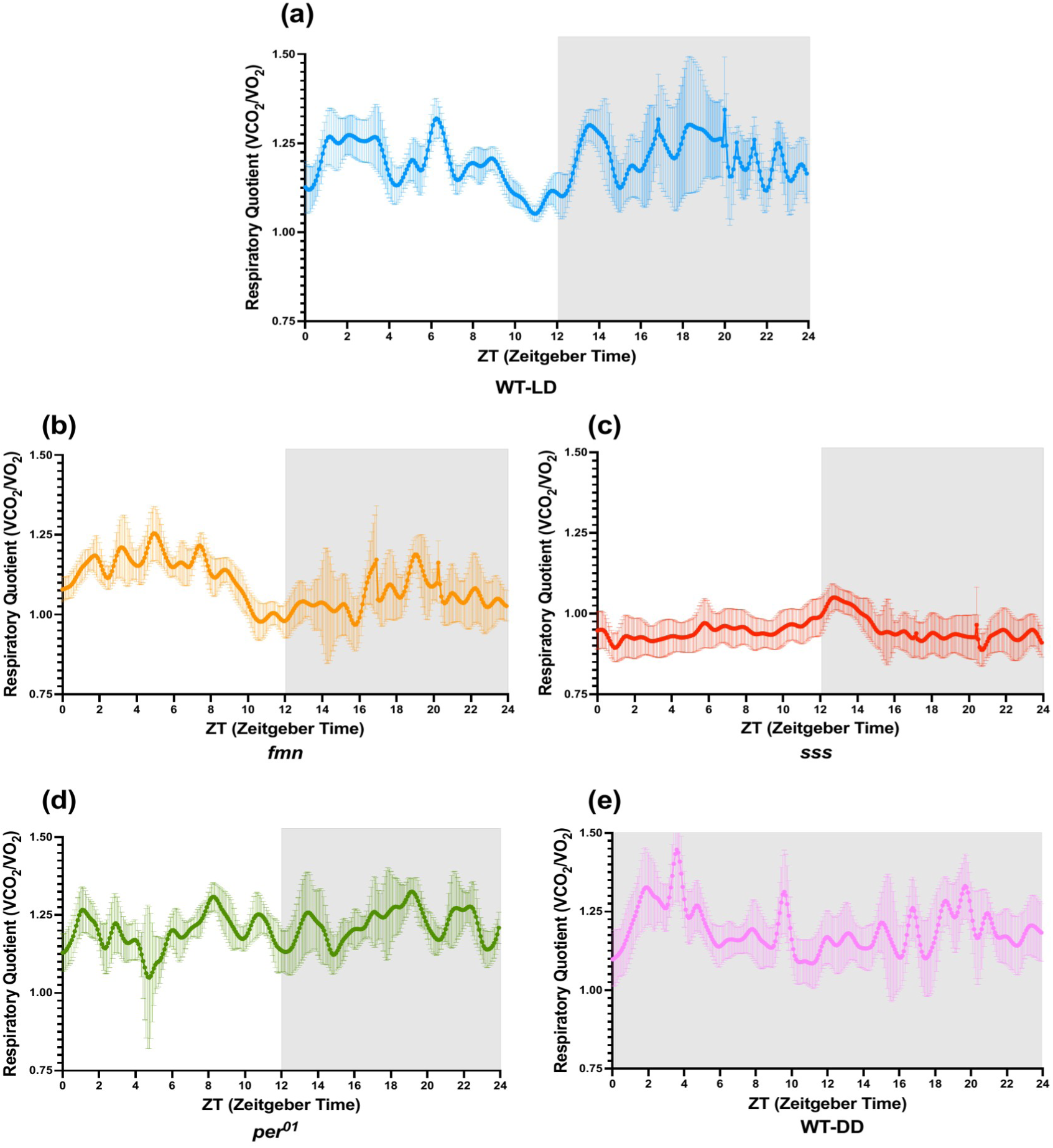
a-e. Respiratory quotient (RQ) profiles across the 24-hour recording period in different genotypes. RQ (VCO₂/VO₂) was continuously recorded at one-second intervals from Zeitgeber Time (ZT) 0 to 24 and subsequently averaged into 5-minute bins for analysis. Traces represent mean RQ values across the 24-hour recording period for wild-type flies under light-dark conditions (WT-LD), short-sleep mutants *fumin* (*fmn*) and *sleepless* (*sss*), the circadian clock mutant *period*^01^ (*per^01^*), and wild-type flies maintained in constant darkness (WT-DD). For all panels, traces are presented as mean ± SEM. Data represent approximately 300 flies per genotype across three experimental days (25 flies/chamber, four chambers/experiment). The chamber was used as the experimental unit; *n* denotes the number of chambers.

**Figure S4.**
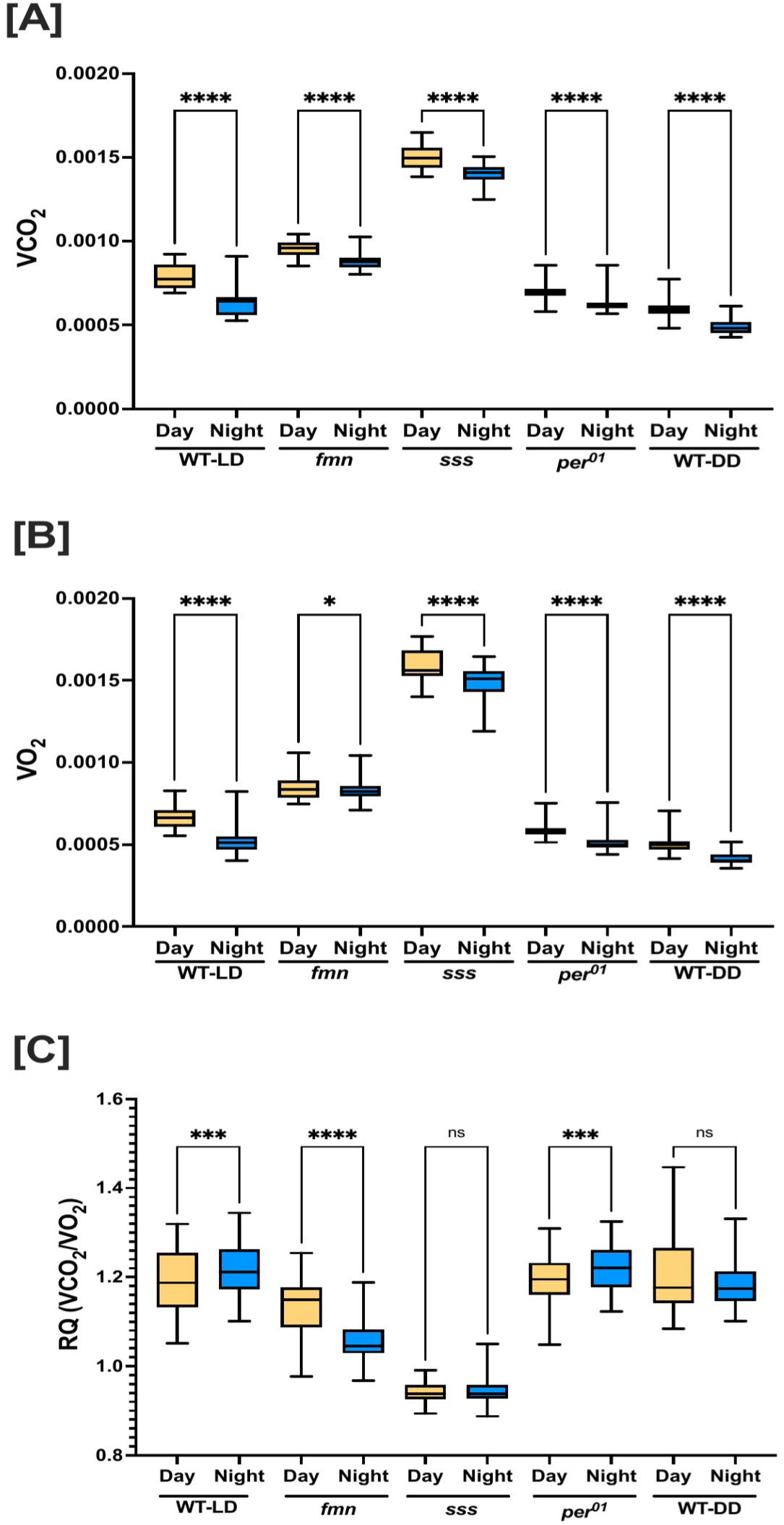
Day-night differences in genotype-specific metabolic rates. Boxplots depict day (ZT0-12) and night (ZT12-24) averages plotted side-by-side for each genotype to facilitate direct day-night comparison. Panels show genotype-specific differences in (A) carbon dioxide production (VCO₂), (B) oxygen consumption (VO₂), and (C) respiratory quotient (RQ). Genotypes include wild-type flies under light-dark conditions (WT-LD), wild-type flies maintained in constant darkness (WT-DD), short-sleep mutants *fumin* (*fmn*) and *sleepless* (*sss*), and the circadian mutant *period^01^* (*per^01^*). Measurements were acquired continuously using a flow-through MAVEn system and binned into day and night intervals. Group differences were assessed using the Kruskal-Wallis test followed by Dunn’s multiple comparisons post hoc test. Significance is denoted as: p < 0.05 (*), p < 0.01 (**), p < 0.001 (***); ns = not significant. Source data are the same continuous respirometry recordings shown in Figure 1. Data represent approximately 300 flies per genotype across three experimental days (25 flies/chamber, four chambers/experiment). The chamber was used as the experimental unit; *n* denotes the number of chambers.

**Supplementary Table 1.**
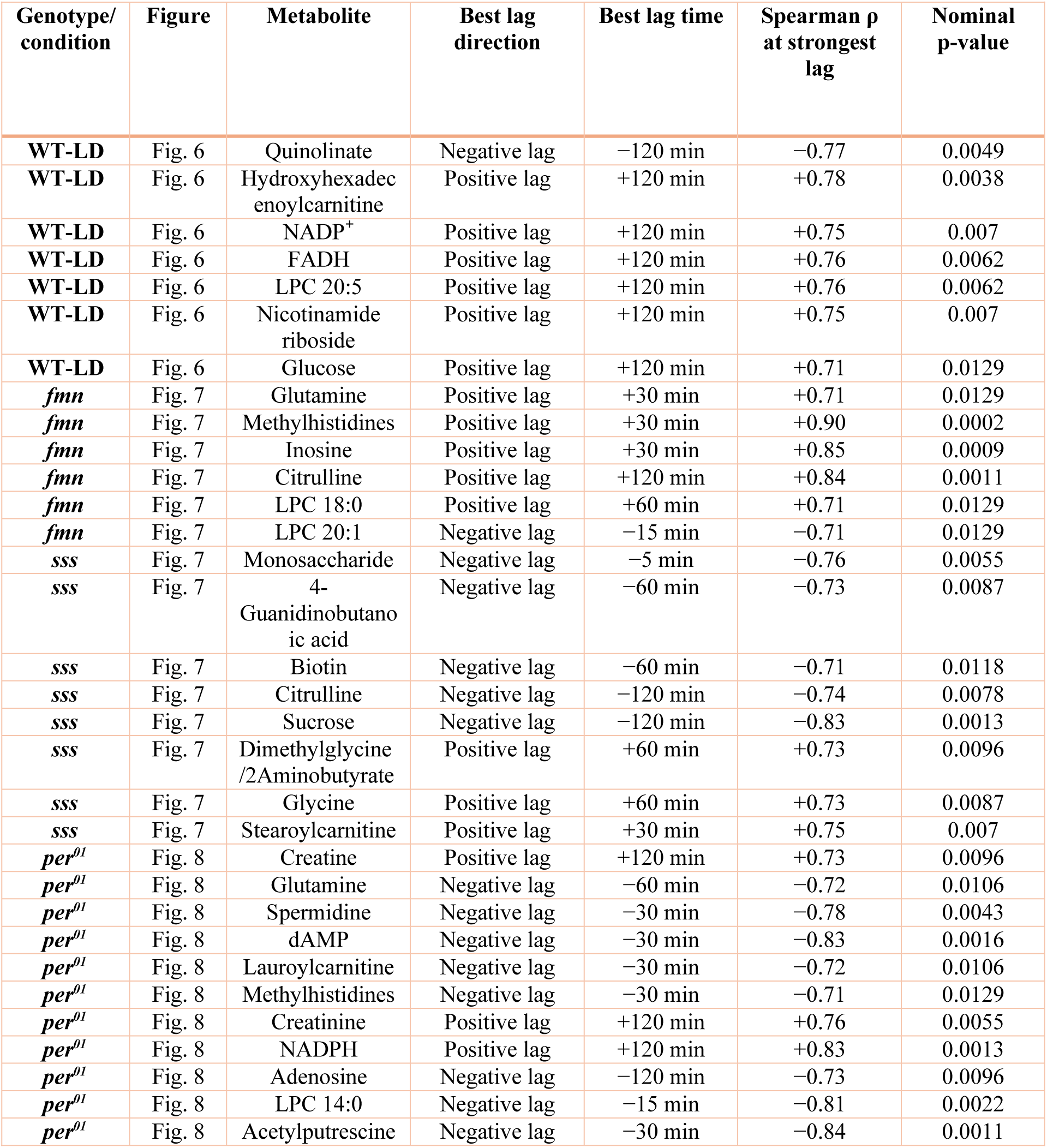
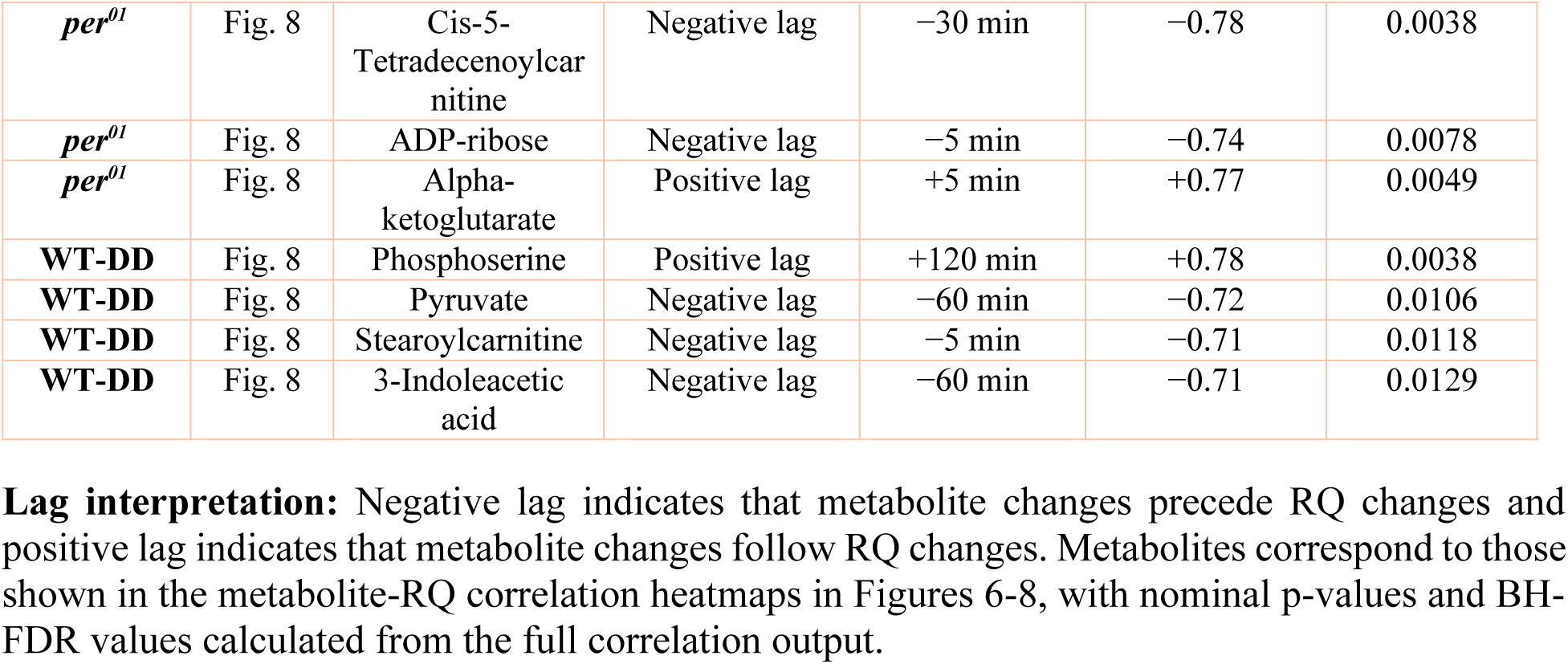
Metabolites showing strong lag-dependent correlations with respiratory quotient (RQ) across genotypes and lighting conditions. Note: Metabolites were included if they showed a strong metabolite–RQ association at one or more temporal lags, defined as |ρ| ≥ 0.7 with nominal p < 0.05. Nominal p-values are reported at the strongest lag for each metabolite. Benjamini–Hochberg false discovery rate (BH-FDR) adjusted p-values were calculated from the full metabolite × lag nominal p-value set within each genotype/condition, including all metabolites and all lag time points tested, and are reported at the strongest lag. BH-FDR adjusted p < 0.05 was considered significant after multiple-testing correction.

| Genotype/<br>condition | Figure | Metabolite | Best lag<br>direction | Best lag time | Spearman $\rho$<br>at strongest<br>lag | Nominal<br>p-value |
| --- | --- | --- | --- | --- | --- | --- |
| WT-LD | Fig. 6 | Quinolate | Negative lag | –120 min | –0.77 | 0.0049 |
| WT-LD | Fig. 6 | Hydroxyhexadec<br>enoylcarnitine | Positive lag | +120 min | +0.78 | 0.0038 |
| WT-LD | Fig. 6 | NADP<sup>+</sup> | Positive lag | +120 min | +0.75 | 0.007 |
| WT-LD | Fig. 6 | FADH | Positive lag | +120 min | +0.76 | 0.0062 |
| WT-LD | Fig. 6 | LPC 20:5 | Positive lag | +120 min | +0.76 | 0.0062 |
| WT-LD | Fig. 6 | Nicotinamide<br>riboside | Positive lag | +120 min | +0.75 | 0.007 |
| WT-LD | Fig. 6 | Glucose | Positive lag | +120 min | +0.71 | 0.0129 |
| *fmn* | Fig. 7 | Glutamine | Positive lag | +30 min | +0.71 | 0.0129 |
| *fmn* | Fig. 7 | Methylhistidines | Positive lag | +30 min | +0.90 | 0.0002 |
| *fmn* | Fig. 7 | Inosine | Positive lag | +30 min | +0.85 | 0.0009 |
| *fmn* | Fig. 7 | Citrulline | Positive lag | +120 min | +0.84 | 0.0011 |
| *fmn* | Fig. 7 | LPC 18:0 | Positive lag | +60 min | +0.71 | 0.0129 |
| *fmn* | Fig. 7 | LPC 20:1 | Negative lag | –15 min | –0.71 | 0.0129 |
| *sss* | Fig. 7 | Monosaccharide | Negative lag | –5 min | –0.76 | 0.0055 |
| *sss* | Fig. 7 | 4-Guanidinobutano<br>ic acid | Negative lag | –60 min | –0.73 | 0.0087 |
| *sss* | Fig. 7 | Biotin | Negative lag | –60 min | –0.71 | 0.0118 |
| *sss* | Fig. 7 | Citrulline | Negative lag | –120 min | –0.74 | 0.0078 |
| *sss* | Fig. 7 | Sucrose | Negative lag | –120 min | –0.83 | 0.0013 |
| *sss* | Fig. 7 | Dimethylglycine<br>/2Aminobutyrate | Positive lag | +60 min | +0.73 | 0.0096 |
| *sss* | Fig. 7 | Glycine | Positive lag | +60 min | +0.73 | 0.0087 |
| *sss* | Fig. 7 | Stearoylcarnitine | Positive lag | +30 min | +0.75 | 0.007 |
| *per*<sup>01</sup> | Fig. 8 | Creatine | Positive lag | +120 min | +0.73 | 0.0096 |
| *per*<sup>01</sup> | Fig. 8 | Glutamine | Negative lag | –60 min | –0.72 | 0.0106 |
| *per*<sup>01</sup> | Fig. 8 | Spermidine | Negative lag | –30 min | –0.78 | 0.0043 |
| *per*<sup>01</sup> | Fig. 8 | dAMP | Negative lag | –30 min | –0.83 | 0.0016 |
| *per*<sup>01</sup> | Fig. 8 | Lauroylcarnitine | Negative lag | –30 min | –0.72 | 0.0106 |
| *per*<sup>01</sup> | Fig. 8 | Methylhistidines | Negative lag | –30 min | –0.71 | 0.0129 |
| *per*<sup>01</sup> | Fig. 8 | Creatinine | Positive lag | +120 min | +0.76 | 0.0055 |
| *per*<sup>01</sup> | Fig. 8 | NADPH | Positive lag | +120 min | +0.83 | 0.0013 |
| *per*<sup>01</sup> | Fig. 8 | Adenosine | Negative lag | –120 min | –0.73 | 0.0096 |
| *per*<sup>01</sup> | Fig. 8 | LPC 14:0 | Negative lag | –15 min | –0.81 | 0.0022 |
| *per*<sup>01</sup> | Fig. 8 | Acetylputrescine | Negative lag | –30 min | –0.84 | 0.0011 |
| *per<sup>01</sup>* | Fig. 8 | Cis-5-Tetradecenoylcarnitine | Negative lag | −30 min | −0.78 | 0.0038 |
| *per<sup>01</sup>* | Fig. 8 | ADP-ribose | Negative lag | −5 min | −0.74 | 0.0078 |
| *per<sup>01</sup>* | Fig. 8 | Alpha-ketoglutarate | Positive lag | +5 min | +0.77 | 0.0049 |
| **WT-DD** | Fig. 8 | Phosphoserine | Positive lag | +120 min | +0.78 | 0.0038 |
| **WT-DD** | Fig. 8 | Pyruvate | Negative lag | −60 min | −0.72 | 0.0106 |
| **WT-DD** | Fig. 8 | Stearoylcarnitine | Negative lag | −5 min | −0.71 | 0.0118 |
| **WT-DD** | Fig. 8 | 3-Indoleacetic acid | Negative lag | −60 min | −0.71 | 0.0129 |

**Supplementary Table 2.**
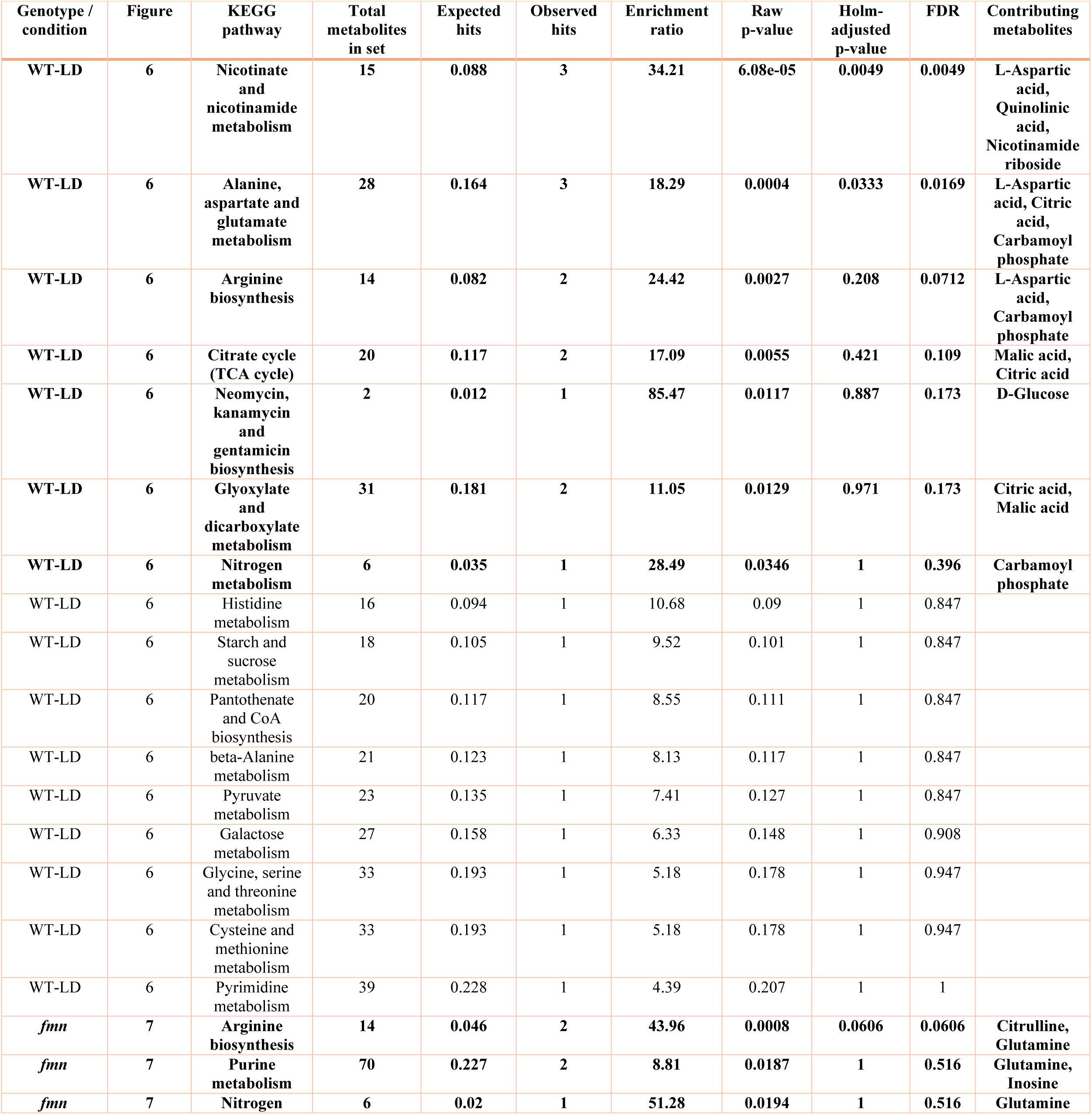
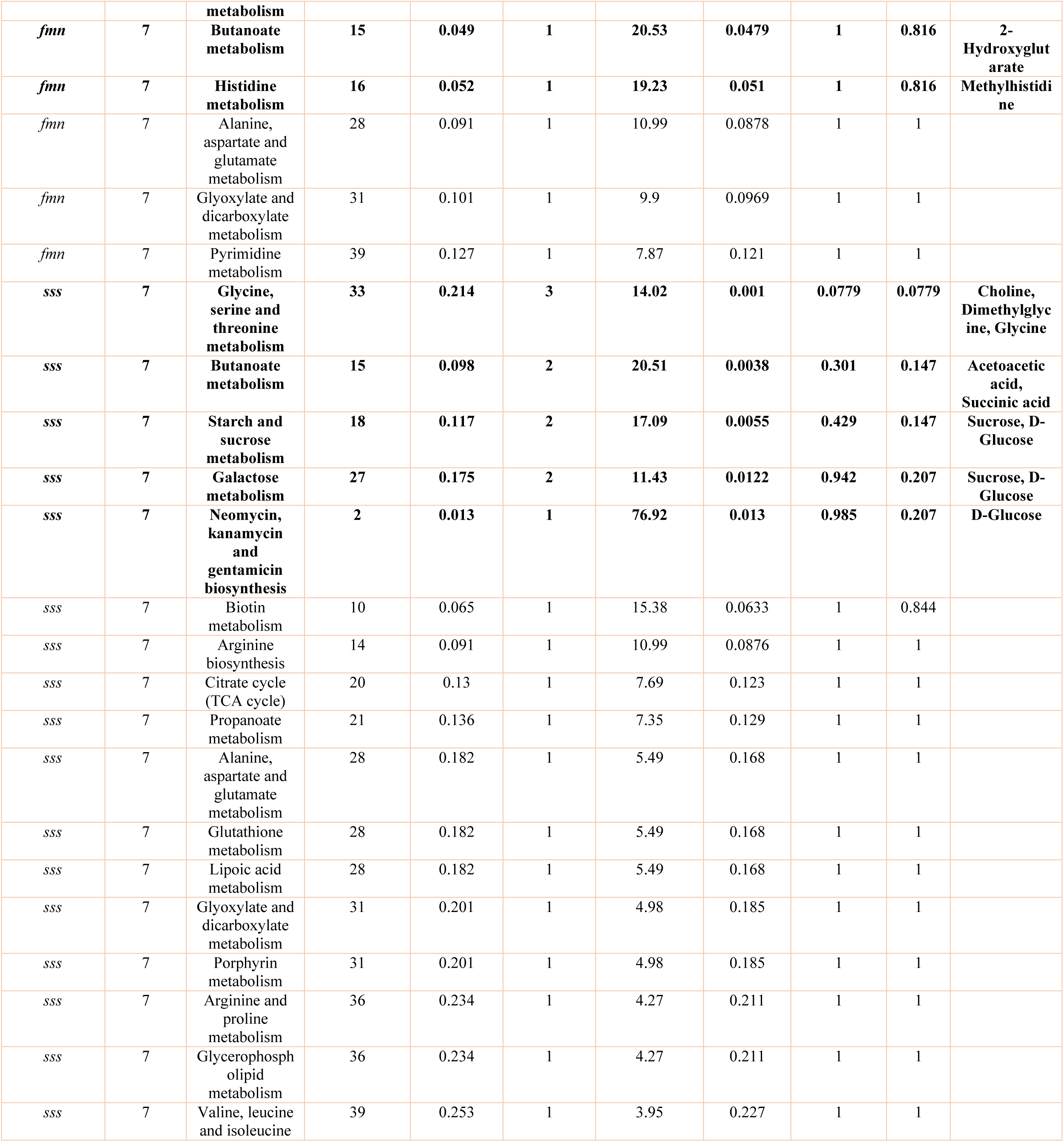
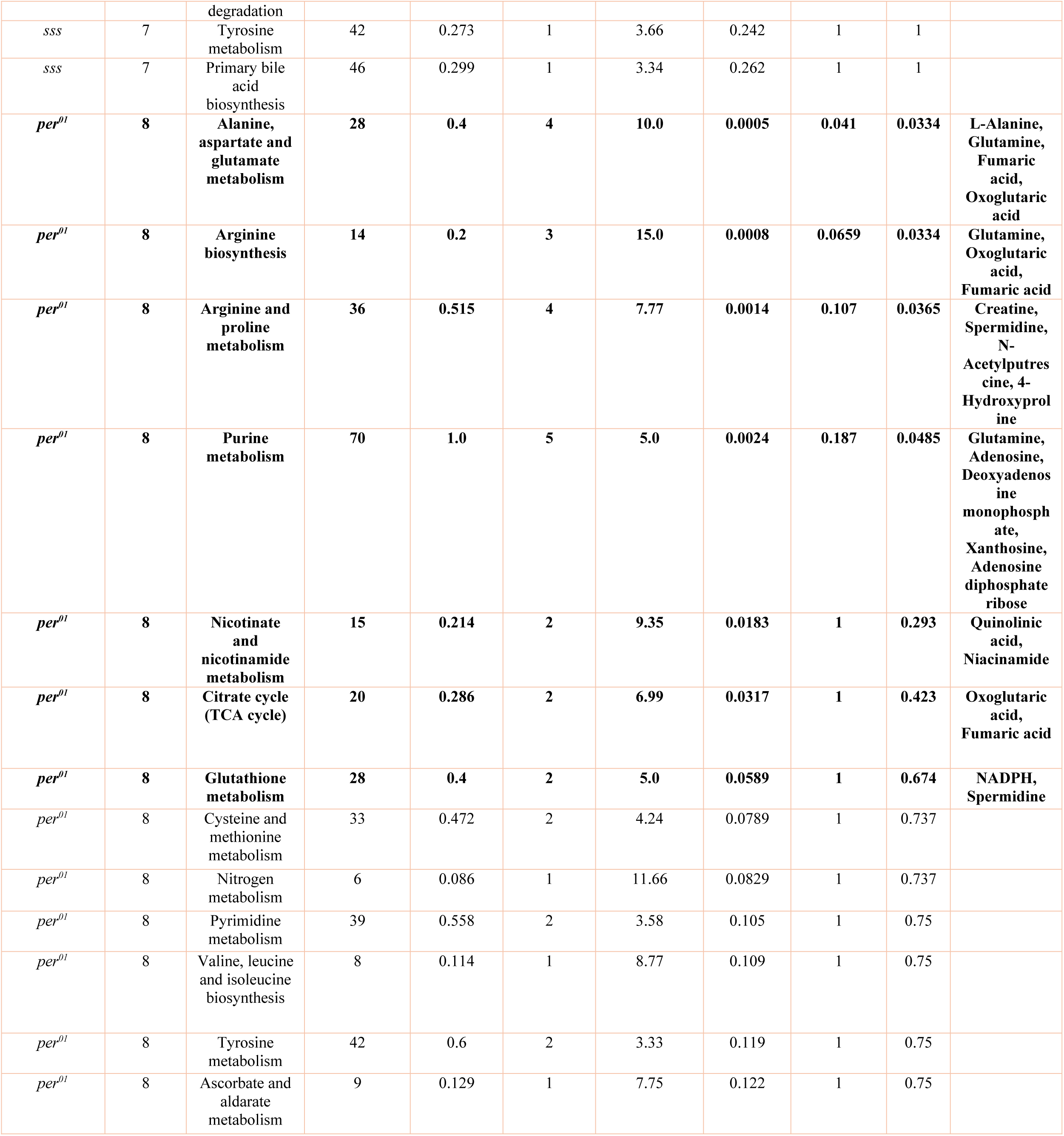
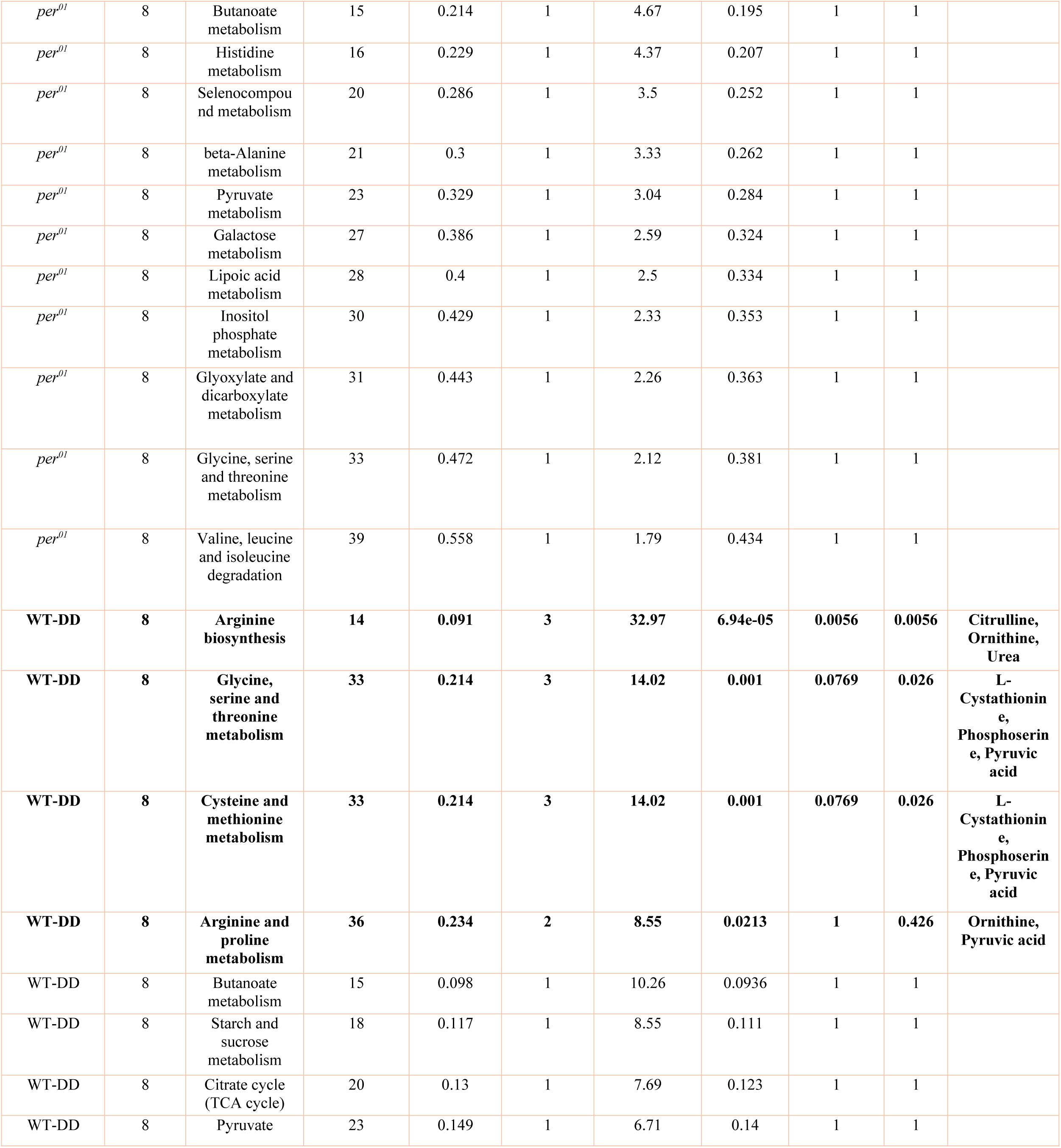
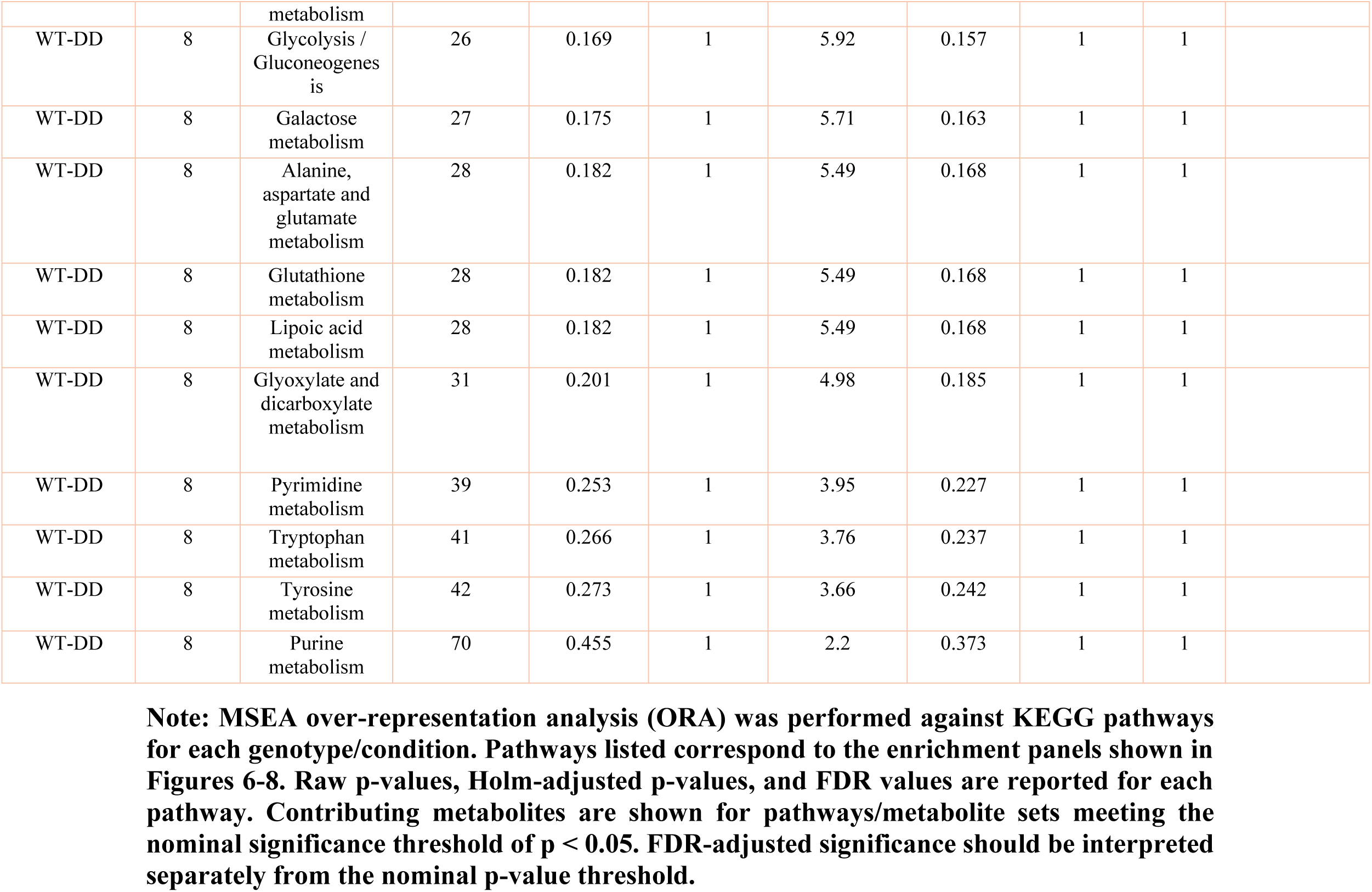
Metabolite Set Enrichment Analysis (MSEA ORA)

**Supplementary Table 3.** Experimental unit, sample size, and error representation for respirometry-based figures. This table summarizes the experimental unit, chamber-level sample size, and error representation for all respirometry-based analyses shown in Figures 1-3 and Figures S1-S4. For all respirometry analyses, the chamber was used as the experimental unit. Each chamber contained 25 flies, and each genotype/condition was measured across four chambers per experiment over three experimental days, corresponding to *n* = 12 chambers per genotype/condition unless otherwise indicated.

| Figure/panel | Data shown | Genotypes /Conditions | Experimental unit | n per genotype/condition | Error bars/shaded bands |
| --- | --- | --- | --- | --- | --- |
| **Fig. 1a-c** | 5-min VCO<sub>2</sub>, VO<sub>2</sub>, RQ traces | WT-LD, *fmn*, *sss*, *per*<sup>01</sup>, WT-DD | Chamber | n = 12 chambers per genotype | Mean ± SEM |
| **Fig. 1d-f** | 24-h average VCO<sub>2</sub>, VO<sub>2</sub>, RQ | WT-LD, *fmn*, *sss*, *per*<sup>01</sup>, WT-DD | Chamber | n = 12 chambers per genotype | Mean ± SEM or boxplot details |
| **Fig. 1g-i** | Body weight / normalized VCO<sub>2</sub>, VO<sub>2</sub> | WT, *fmn*, *sss*, *per*<sup>01</sup> | Chamber | n = 12 chambers per genotype | Mean ± SEM |
| **Fig. 2** | Rhythmicity time course | WT-LD, *fmn*, *sss*, *per*<sup>01</sup>, WT-DD | Chamber | n = 12 chambers per genotype | Mean ± SEM |
| **Fig. 3** | Phase/peak timing | WT-LD, *fmn*, *sss*, *per*<sup>01</sup>, WT-DD | Chamber | n = 12 chambers per genotype | Not applicable / phase estimate |
| **Fig. S1-S3** | Individual VCO<sub>2</sub>, VO<sub>2</sub>, RQ traces | WT-LD, *fmn*, *sss*, *per*<sup>01</sup>, WT-DD | Chamber | 12 chambers per genotype | Mean ± SEM |
| **Fig. S4** | Day vs night averages | WT-LD, *fmn*, *sss*, *per*<sup>01</sup>, WT-DD | Chamber | 12 chambers per genotype | Mean ± SEM or boxplot details |

**Note: Chamber was used as the experimental unit for all respirometry analyses. Each chamber contained 25 flies, and each genotype/condition was measured across four chambers per experiment over three experimental days, corresponding to $n = 12$ chambers per genotype/condition unless otherwise indicated. Error bars or shaded bands represent SEM. For boxplots, the plotted distribution/error representation is indicated in the corresponding figure legend.**

## REFERENCES

1. Bass, J. and J.S. Takahashi, Circadian integration of metabolism and energetics. Science, 2010. 330(6009): p. 1349–54.

2. Panda, S., Circadian physiology of metabolism. Science, 2016. 354(6315): p. 1008–1015.

3. Sharma, S. and M. Kavuru, Sleep and metabolism: an overview. Int J Endocrinol, 2010. 2010.

4. Aurora, R.N. and N.M. Punjabi, Obstructive sleep apnoea and type 2 diabetes mellitus: a bidirectional association. Lancet Respir Med, 2013. 1(4): p. 329–38.

5. Leproult, R. and E. Van Cauter, Role of sleep and sleep loss in hormonal release and metabolism. Endocr Dev, 2010. 17: p. 11–21.

6. Reutrakul, S. and E. Van Cauter, Interactions between sleep, circadian function, and glucose metabolism: implications for risk and severity of diabetes. Ann N Y Acad Sci, 2014. 1311: p. 151–73.

7. Sulli, G., et al., Training the Circadian Clock, Clocking the Drugs, and Drugging the Clock to Prevent, Manage, and Treat Chronic Diseases. Trends Pharmacol Sci, 2018. 39(9): p. 812–827.

8. Hendricks, J.C., et al., Rest in Drosophila is a sleep-like state. Neuron, 2000. 25(1): p. 129–38.

9. Allada, R. and J.M. Siegel, Unearthing the phylogenetic roots of sleep. Curr Biol, 2008. 18(15): p. R670–R679.

10. Donelson, N.C., et al., High-resolution positional tracking for long-term analysis of Drosophila sleep and locomotion using the "tracker" program. PLoS One, 2012. 7(5): p. e37250.

11. Speakman, J.R., Measuring energy metabolism in the mouse - theoretical, practical, and analytical considerations. Front Physiol, 2013. 4: p. 34.

12. Even, P.C., A. Mokhtarian, and A. Pele, Practical aspects of indirect calorimetry in laboratory animals. Neurosci Biobehav Rev, 1994. 18(3): p. 435–47.

13. McClave, S.A. and H.L. Snider, Use of indirect calorimetry in clinical nutrition. Nutr Clin Pract, 1992. 7(5): p. 207–21.

14. Frayn, K.N., Calculation of substrate oxidation rates in vivo from gaseous exchange. J Appl Physiol Respir Environ Exerc Physiol, 1983. 55(2): p. 628–34.

15. Arch, J.R., et al., Some mathematical and technical issues in the measurement and interpretation of open-circuit indirect calorimetry in small animals. Int J Obes (Lond), 2006. 30(9): p. 1322–31.

16. Yatsenko, A.S., et al., Measurement of metabolic rate in Drosophila using respirometry. J Vis Exp, 2014(88): p. e51681.

17. Wiggin, T.D., et al., Covert sleep-related biological processes are revealed by probabilistic analysis in Drosophila. Proc Natl Acad Sci U S A, 2020. 117(18): p. 10024–10034.

18. Brown, E.B., J. Klok, and A.C. Keene, Measuring metabolic rate in single flies during sleep and waking states via indirect calorimetry. J Neurosci Methods, 2022. 376: p. 109606.

19. Cedernaes, J., H.B. Schioth, and C. Benedict, Determinants of shortened, disrupted, and mistimed sleep and associated metabolic health consequences in healthy humans. Diabetes, 2015. 64(4): p. 1073–80.

20. Katayose, Y., et al., Metabolic rate and fuel utilization during sleep assessed by whole-body indirect calorimetry. Metabolism, 2009. 58(7): p. 920–6.

21. Kume, K., et al., Dopamine is a regulator of arousal in the fruit fly. J Neurosci, 2005. 25(32): p. 7377–84.

22. Koh, K., et al., Identification of SLEEPLESS, a sleep-promoting factor. Science, 2008. 321(5887): p. 372–6.

23. Chen, W.F., et al., A neuron-glia interaction involving GABA transaminase contributes to sleep loss in sleepless mutants. Mol Psychiatry, 2015. 20(2): p. 240–51.

24. Wu, M.N., et al., SLEEPLESS, a Ly-6/neurotoxin family member, regulates the levels, localization and activity of Shaker. Nat Neurosci, 2010. 13(1): p. 69–75.

25. Konopka, R.J. and S. Benzer, Clock mutants of Drosophila melanogaster. Proc Natl Acad Sci U S A, 1971. 68(9): p. 2112–6.

26. Stahl, B.A., et al., Sleep-Dependent Modulation of Metabolic Rate in Drosophila. Sleep, 2017. 40(8).

27. Malik, D.M., et al., Altered Metabolism during the Dark Period in Drosophila Short Sleep Mutants. J Proteome Res, 2024. 23(9): p. 3823–3836.

28. Yuan, M., et al., A positive/negative ion-switching, targeted mass spectrometry-based metabolomics platform for bodily fluids, cells, and fresh and fixed tissue. Nat Protoc, 2012. 7(5): p. 872–81.

29. Rhoades, S.D., et al., Circadian- and Light-driven Metabolic Rhythms in Drosophila melanogaster. J Biol Rhythms, 2018. 33(2): p. 126–136.

30. Lighton, J.R.B., Measuring Metabolic Rates: A Manual for Scientists. 2008.

31. Malik, D.M., S. Rhoades, and A. Weljie, Extraction and Analysis of Pan-metabolome Polar Metabolites by Ultra Performance Liquid Chromatography-Tandem Mass Spectrometry (UPLC-MS/MS). Bio Protoc, 2018. 8(3).

32. Sengupta, A. and A.M. Weljie, NMR Spectroscopy-Based Metabolic Profiling of Biospecimens. Curr Protoc Protein Sci, 2019. 98(1): p. e98.

33. Brooks, T.G., et al., Nitecap: An Exploratory Circadian Analysis Web Application. J Biol Rhythms, 2022. 37(1): p. 43–52.

34. Thaben, P.F. and P.O. Westermark, Detecting rhythms in time series with RAIN. J Biol Rhythms, 2014. 29(6): p. 391–400.

35. Hughes, M.E., J.B. Hogenesch, and K. Kornacker, JTK_CYCLE: an efficient nonparametric algorithm for detecting rhythmic components in genome-scale data sets. J Biol Rhythms, 2010. 25(5): p. 372–80.

36. Tennessen, J.M., et al., Methods for studying metabolism in Drosophila. Methods, 2014. 68(1): p. 105–15.

37. Cirelli, C., The genetic and molecular regulation of sleep: from fruit flies to humans. Nat Rev Neurosci, 2009. 10(8): p. 549–60.

38. Johnson, C.H., J. Ivanisevic, and G. Siuzdak, Metabolomics: beyond biomarkers and towards mechanisms. Nat Rev Mol Cell Biol, 2016. 17(7): p. 451–9.

39. Patti, G.J., O. Yanes, and G. Siuzdak, Innovation: Metabolomics: the apogee of the omics trilogy. Nat Rev Mol Cell Biol, 2012. 13(4): p. 263–9.

40. Wishart, D.S., Emerging applications of metabolomics in drug discovery and precision medicine. Nat Rev Drug Discov, 2016. 15(7): p. 473–84.

41. Wu, M.N., et al., A genetic screen for sleep and circadian mutants reveals mechanisms underlying regulation of sleep in Drosophila. Sleep, 2008. 31(4): p. 465–72.

42. Shi, M., et al., Identification of Redeye, a new sleep-regulating protein whose expression is modulated by sleep amount. Elife, 2014. 3: p. e01473.

43. Ho, J.M., R.P. Barf, and M.R. Opp, Effects of sleep disruption and high fat intake on glucose metabolism in mice. Psychoneuroendocrinology, 2016. 68: p. 47–56.

44. Jung, C.M., et al., Energy expenditure during sleep, sleep deprivation and sleep following sleep deprivation in adult humans. J Physiol, 2011. 589(Pt 1): p. 235–44.

45. Woller, A. and D. Gonze, Circadian Misalignment and Metabolic Disorders: A Story of Twisted Clocks. Biology (Basel), 2021. 10(3).

46. Dibner, C. and U. Schibler, Circadian timing of metabolism in animal models and humans. J Intern Med, 2015. 277(5): p. 513–27.

47. O’Neill, J.S. and A.B. Reddy, Circadian clocks in human red blood cells. Nature, 2011. 469(7331): p. 498–503.

48. Nakahata, Y., et al., Circadian control of the NAD+ salvage pathway by CLOCK-SIRT1. Science, 2009. 324(5927): p. 654–7.

49. Green, C.B., J.S. Takahashi, and J. Bass, The meter of metabolism. Cell, 2008. 134(5): p. 728–42.

50. Rey, G., et al., Metabolic oscillations on the circadian time scale in Drosophila cells lacking clock genes. Mol Syst Biol, 2018. 14(8): p. e8376.

51. Peek, C.B., et al., Circadian clock NAD+ cycle drives mitochondrial oxidative metabolism in mice. Science, 2013. 342(6158): p. 1243417.

52. Sharon, G., et al., Commensal bacteria play a role in mating preference of Drosophila melanogaster. Proc Natl Acad Sci U S A, 2010. 107(46): p. 20051–6.

53. Wong, A.C., A.J. Dobson, and A.E. Douglas, Gut microbiota dictates the metabolic response of Drosophila to diet. J Exp Biol, 2014. 217(Pt 11): p. 1894–901.

54. Stenvers, D.J., et al., Circadian clocks and insulin resistance. Nat Rev Endocrinol, 2019. 15(2): p. 75–89.

55. Leone, V., et al., Effects of diurnal variation of gut microbes and high-fat feeding on host circadian clock function and metabolism. Cell Host Microbe, 2015. 17(5): p. 681–9.

56. Donlea, J.M., D. Pimentel, and G. Miesenbock, Neuronal Machinery of Sleep Homeostasis in Drosophila. Neuron, 2014. 81(6): p. 1442.

57. Liu, S., et al., Sleep Drive Is Encoded by Neural Plastic Changes in a Dedicated Circuit. Cell, 2016. 165(6): p. 1347–1360.

58. Brosnan, J.T., Glutamate, at the interface between amino acid and carbohydrate metabolism. J Nutr, 2000. 130(4S Suppl): p. 988S–90S.

59. Reinecke, F., J.A. Smeitink, and F.H. van der Westhuizen, OXPHOS gene expression and control in mitochondrial disorders. Biochim Biophys Acta, 2009. 1792(12): p. 1113–21.

60. Vaccaro, A., et al., Sleep Loss Can Cause Death through Accumulation of Reactive Oxygen Species in the Gut. Cell, 2020. 181(6): p. 1307–1328 e15.

61. Mentch, S.J., et al., Histone Methylation Dynamics and Gene Regulation Occur through the Sensing of One-Carbon Metabolism. Cell Metab, 2015. 22(5): p. 861–73.

62. Locasale, J.W., Serine, glycine and one-carbon units: cancer metabolism in full circle. Nat Rev Cancer, 2013. 13(8): p. 572–83.

63. Owen, O.E., S.C. Kalhan, and R.W. Hanson, The key role of anaplerosis and cataplerosis for citric acid cycle function. J Biol Chem, 2002. 277(34): p. 30409–12.

64. Finkel, T., et al., The ins and outs of mitochondrial calcium. Circ Res, 2015. 116(11): p. 1810–9.

65. Keene, A.C., et al., Clock and cycle limit starvation-induced sleep loss in Drosophila. Curr Biol, 2010. 20(13): p. 1209–15.

66. Hatori, M., et al., Time-restricted feeding without reducing caloric intake prevents metabolic diseases in mice fed a high-fat diet. Cell Metab, 2012. 15(6): p. 848–60.

67. Hardin, P.E., The circadian timekeeping system of Drosophila. Curr Biol, 2005. 15(17): p. R714–22.

68. Wilking, M., et al., Circadian rhythm connections to oxidative stress: implications for human health. Antioxid Redox Signal, 2013. 19(2): p. 192–208.

69. Kriebs, A., et al., Circadian repressors CRY1 and CRY2 broadly interact with nuclear receptors and modulate transcriptional activity. Proc Natl Acad Sci U S A, 2017. 114(33): p. 8776–8781.

70. Allada, R. and B.Y. Chung, Circadian organization of behavior and physiology in Drosophila. Annu Rev Physiol, 2010. 72: p. 605–24.

71. Dubowy, C. and A. Sehgal, Circadian Rhythms and Sleep in Drosophila melanogaster. Genetics, 2017. 205(4): p. 1373–1397.

72. Dallmann, R., S.A. Brown, and F. Gachon, Chronopharmacology: new insights and therapeutic implications. Annu Rev Pharmacol Toxicol, 2014. 54: p. 339–61.

73. Mentch, S.J. and J.W. Locasale, One-carbon metabolism and epigenetics: understanding the specificity. Ann N Y Acad Sci, 2016. 1363(1): p. 91–8.

74. Hughes, M.E., et al., Harmonics of circadian gene transcription in mammals. PLoS Genet, 2009. 5(4): p. e1000442.

75. Malik, D.M., et al., Glucose is dynamically regulated by time of day in humans and Drosophila. PLoS Biol, 2026. 24(4): p. e3003717.

